# A marginal habitat, but not a sink: Ecological genetics reveal a diversification hotspot for marine invertebrates in the brackish Baltic Sea

**DOI:** 10.1101/2021.06.06.447071

**Authors:** Jonas C. Geburzi, Nele Heuer, Lena Homberger, Jana Kabus, Zoe Moesges, Kira Ovenbeck, Dirk Brandis, Christine Ewers-Saucedo

## Abstract

**Aim:** Environmental gradients have emerged as important barriers structuring populations and species distributions. We set out to test whether a strong salinity gradient from marine to brackish, represented in a marginal northern European sea, should be considered a diversification hotspot or a population sink, and to identify life history traits that correlate with either evolutionary trajectory.

**Location:** The Baltic Sea, the North Sea and their transition zone.

**Methods:** We accumulated mitochondrial cytochrome oxidase subunit 1 sequence data and data on the distribution, salinity tolerance and life history for 28 species belonging to the Cnidaria, Crustacea, Echinodermata, Mollusca, Polychaeta and Gastrotricha, including seven non-native species. We calculated measures of genetic diversity and differentiation across the environmental gradient, coalescent times and migration rates between North and Baltic Sea populations, and analysed correlations between genetic and life history data.

**Results:** The majority of investigated species is either genetically differentiated and/or is adapted to the lower salinity conditions of the Baltic Sea. Moreover, the species exhibiting population structure have a range of patterns of genetic diversity in comparison to the North Sea, from lower in the Baltic Sea to higher in the Baltic Sea, or equally diverse in North and Baltic Sea.

**Main conclusions:** Our results indicate that the Baltic Sea should be considered a diversification hotspot: The diversity of genetic patterns points towards independent trajectories in the Baltic compared to the North Sea. At the same time, we found limited evidence for the traditional scenario of the Baltic Sea as a population sink with lower diversity and strong gene flow. The North Sea - Baltic Sea region provides a unique setting to study evolutionary adaptation during colonization processes at different stages by jointly considering native and non-native species.

## Introduction

Environmental gradients have emerged as important barriers, structuring populations and species distributions. The environment may be particularly important in the marine realm, where impenetrable barriers, such as land masses, are relatively rare (Blanco-Bercial et al., 2011; Ewers-Saucedo and Wares, 2020). In particular, temperature (Ewers-Saucedo et al., 2016), salinity (Sjöqvist et al., 2015) and water depth (Prada and Hellberg, 2021) may result in differentially adapted populations with limited gene flow. In addition to adaptive limits to gene flow, several barriers to dispersal exist. Currents and upwelling limit dispersal for benthic invertebrates with a planktonic larval phase, albeit these barriers are not universal (Wares et al., 2001; Kelly and Palumbi, 2010; Haye et al., 2014). Just as important are stretches of unsuitable habitat, for example long sandy beaches for rocky shore specialists (Ayre et al., 2009; Wares, 2019). In lieu of a planktonic phase or other long-distance dispersal mechanisms, small-scale population structure is commonplace in benthic species (Palumbi, 1994; Kyle and Boulding, 2000; Haye et al., 2014; Ewers-Saucedo and Wares, 2020).

A marine region characterized by both restricted water movement and strong environmental gradients is the North Sea - Baltic Sea region. The Baltic Sea is the world’s largest inland brackish water body with a west-to-east salinity gradient. The North Sea connects to the Baltic Sea via the narrow channels of Kattegat, Skagerrak, and the Belt Sea, which is littered with islands and bridges (Fig. 1). Most marine organisms likely colonized the Baltic from the North Sea over the course of the past 8000 years when the Baltic Sea turned from freshwater to brackish after the last glacial maximum (15,000 ya). During the preceding glaciation, the Baltic Sea was covered in ice. Before that, until about 200,000 ya, the geographic region of the Baltic Sea was no sea at all, but a landmass with a large river system (André et al., 2011). Numerous non-native species from other regions of the world have entered the Baltic Sea since, e.g. the Black Sea lineage of the shrimp *Palaemon elegans*, the crab *Hemigrapsus takanoi* from Japan and the clam *Mya arenaria* from North America (Petersen et al., 1992; Behrends et al., 2005; Reuschel et al., 2010; Geburzi et al., 2015). A few species, such as the blue mussel *Mytilus trossulus*, may have colonized the Baltic Sea from the White or Barents Sea during brief periods when it was connected to the Baltic Sea (Väinölä and Strelkov, 2011).

**Figure 1.**
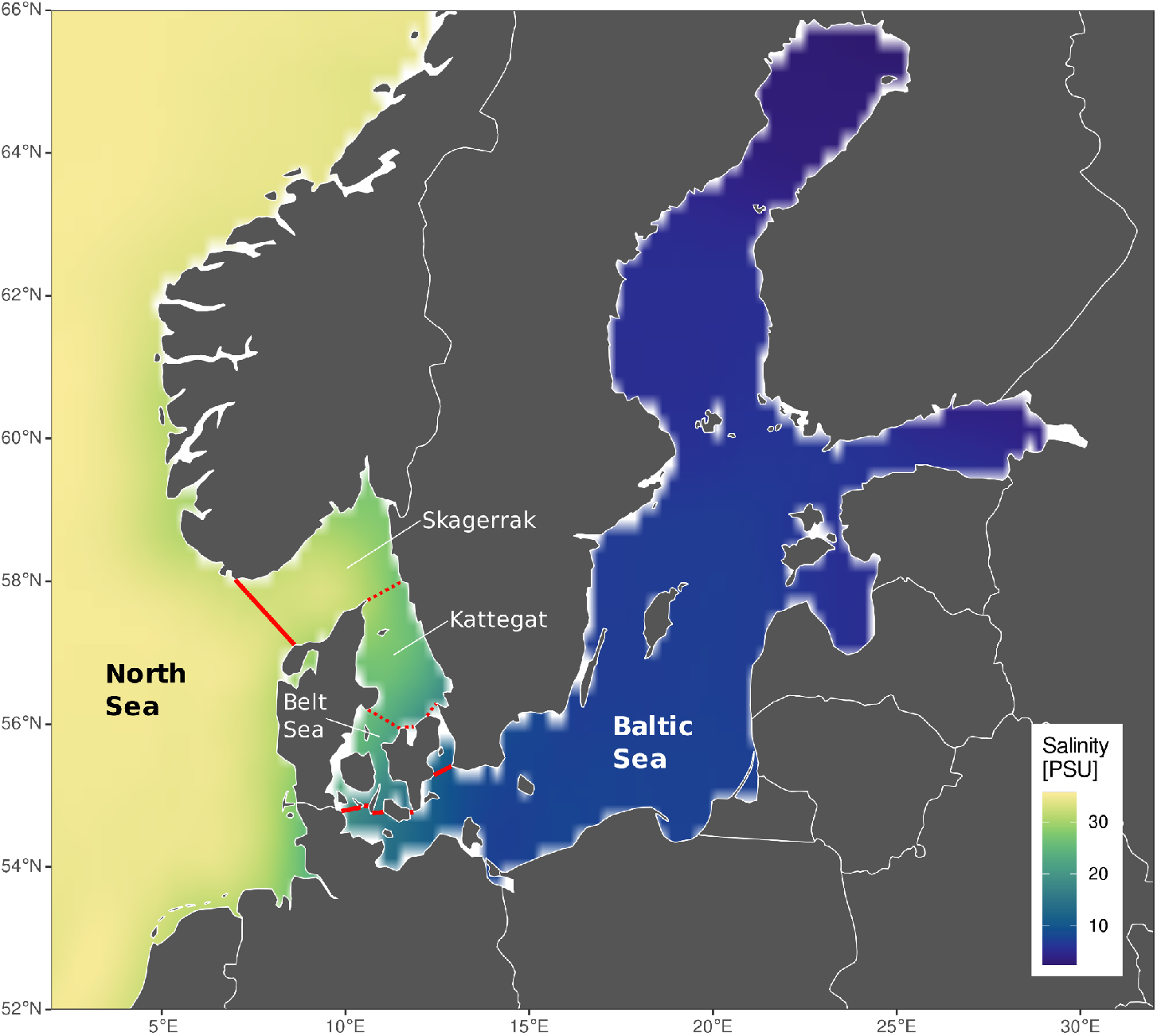
Map of the North Sea - Baltic Sea salinity gradient, showing the decadal interpolated average salinity from 2006-2015. Skagerrak, Kattegat and Belt Sea comprise the transition zone between the North and Baltic Seas, i.e. the area between the solid red lines. Salinity data from Hinrichs & Gouretski (2018) available at https://icdc.cen.uni-hamburg.de/bnsc-project.html.

Despite the young evolutionary age of the Baltic Sea, at least two species evolved in the Baltic Sea, the brown algae *Fucus radicans* (Pereyra et al., 2009), and the Baltic flounder *Platichthys solemdali* (Momigliano et al., 2017). In both cases, adaptation to the salinity gradient has been invoked as the driving force of speciation. Adaptation may also play a role during ecological differentiation between populations: at least five fish species in the Baltic Sea are adapted to low salinity (Larsen et al., 2008; Andersen et al., 2009; Gaggiotti et al., 2009; Papakostas et al., 2012; Berg et al., 2015; Guo et al., 2015), as are the marine amphipods *Gammarus locusta* and *G. oceanicus* (Segerstråle, 1947; den Hartog, 1964; Fenchel and Kolding, 1979), and the jellyfish *Aurelia aurita* has differing reproductive cycles in different parts of the North and Baltic Sea system (Lucas, 2001). The marine diatom *Skeletonema marinoi* shows adaptive growth optima under North or Baltic Sea salinities, as well as genetic differentiation (Sjöqvist et al., 2015).

These findings may represent a general scheme for the Baltic Sea as a diversification hotspot. Corroborating evidence comes from a number of population genetic studies that found significant genetic differentiation across the North Sea - Baltic Sea environmental gradient (Johannesson and André, 2006; Wennerström et al., 2013, 2017). However, other species are not significantly differentiated (Johannesson and André, 2006). These species do not necessarily contradict the diversification hotspot idea. Instead, they may have colonized the Baltic Sea relatively late, not leaving enough time for observable genetic differences to arise. What constitutes “enough time” depends on a species’ demography: large populations need longer to differentiate, as do species with a long generation time (Kingman, 1982). Even little gene flow, which we may expect based on intermittent saltwater inflow from the North Sea, slows down differentiation processes (Kimura and Maruyama, 1971).

Alternatively, the confirmed cases of rapid adaptation in the Baltic Sea could be exceptions to the traditional view of the Baltic Sea fauna as a depauperate branch of the North Sea fauna (Meyer and Möbius, 1865). This idea implies that the Baltic Sea is a population sink, which is supplied with a limited number of propagules from the source population in the North Sea. This hypothesis stems from the observation that the Baltic Sea harbors only a small percentage of the marine species present in the North Sea (Zettler et al., 2018), and that such biogeographic patterns are generally mirrored by phylogeography (Vellend, 2003; Vellend and Geber, 2005; Díaz-Ferguson et al., 2010 for references). In congruence with this argument, most investigated species to date had lower genetic diversity in the Baltic (Johannesson and André, 2006).

It seems plausible that both evolutionary trajectories co-exist, and the path each species takes depends on their life history and demography (Ewers-Saucedo and Wares, 2020). Limited water exchange should influence species with a planktonic phase the most, while species with little dispersal ability might be fastest to adapt locally to environmental conditions (Schluter, 2000; Kisdi, 2002). Moreover, intrinsic environmental tolerances differ between species, so that some species perceive an environmental barrier where others do not. This means that for some species, the entrance of the Baltic Sea may form a significant barrier to gene flow, while other species may cross into the Baltic unhindered.

We set out to test whether the Baltic Sea should be considered a diversification hotspot or a population sink, and to identify life history traits that correlate with either evolutionary trajectory. We analysed newly generated and publicly available genetic barcoding data, and compiled data on life history traits and basin-specific salinity tolerances. Differing salinity tolerances between the two basins point to rapid adaptation, especially in concert with limited gene flow between North and Baltic Sea. Given the diversity in life histories and population sizes, we focused this study on marine invertebrate species, and conducted population genetic simulations to understand the expected outcomes based on limited sampling and the evolutionary young age of the Baltic Sea.

## Methods

### Acquisition of life history data

We searched the literature for information on pelagic larval duration (PLD), adult dispersal ability, adult habitat and salinity tolerance for all investigated species. We included both experimental and observational estimates of salinity tolerance, and discriminated between estimates for populations from within and outside the Baltic Sea where possible. As a further proxy for salinity tolerance, we recorded the most eastern longitude at which a species was reported consistently in the Baltic Sea, based on the OBIS (Ocean Biodiversity Information System, https://obis.org) and GBIF (Global Biodiversity Information Facility, www.gbif.org) databases, as well as distribution records in the literature. For non-native species, we also searched for the year of their first records in the North and Baltic Seas, respectively. A list of the data sources is found in Appendix 1.

### COI sequencing

Over the past ten years, bachelor, masters and doctoral students at the Zoological Museum in Kiel have investigated population genetic differences between North and Baltic Sea populations for a number of marine invertebrates. For each species, the students extracted DNA from a maximum of 20 specimens each from North Sea and Baltic Sea using commercial DNA extraction kits (Roth, Stratec Molecular) or the Chelex method (Walsh et al., 1991). Student-and species-specific information on the respective extraction protocol, primers and PCR settings are available in the Supporting Information 1. The accession numbers for these new sequence data on NCBI GenBank (www.ncbi.nlm.nih.gov) are available in the Supporting Information 2 (column ‘New GenBank Acc.’).

### Acquisition of genetic data

We searched for publicly available cytochrome oxidase subunit 1 (COI) sequence data of species from the western or central Baltic Sea as well as the North Sea. Sequence data for the transition zone (as described in Fig. 1) was generally rare, and we did not include it into our overall analyses, but utilized it to assess the location of genetic breaks where appropriate. We began by extracting data from the studies cited in Johannesson and André (2006). To find newer publicly available sequence data especially for the Baltic Sea, we searched for articles citing Johanneson and André (2006), searched NCBI GenBank for “Baltic Sea’’ and “cytochrome oxidase”, and searched google scholar for “Baltic Sea phylogeography marine” and “Baltic Sea marine population”. A list of the sequence data sources is found in Appendix 2. We downloaded sequence or haplotype data from NCBI GenBank, supplements of publications or the Barcoding of Life Database website (www.boldsystems.org). For accession numbers for these downloaded data from both NCBI GenBank and the Barcoding of Life Database, see Tab. S2. When the sequence data represented haplotypes, rather than sequences for each sampled individual, we reconstructed haplotype frequencies from information within the respective publication.

### Data quality control

We excluded species that had not colonized the Baltic Sea directly via the North Sea or vice versa. However, we kept species where the colonization may have proceeded from the Baltic Sea to the North Sea, as in some brackish water species. We removed highly divergent sequences from cryptic or misidentified species. For each species, we aligned all COI sequences in Geneious v.9.1.8 (Kearse et al., 2012) with the “Map to reference” function, using the longest sequence as reference. This appeared to be a faster, more reliable alignment approach than aligning with a “Multiple align” algorithm. We checked the alignment for gaps, removed short sequences, and trimmed all remaining sequences to the same length. This means that different species have different alignment lengths. We removed species with a final alignment length below 400bp, as shorter sequences are likely to harbor less genetic diversity, and thus may lead to underestimates of diversity and differentiation.

### Population genetic analyses

All analyses were conducted in the R environment (R Core Team, 2019). For each species, we reconstructed haplotype networks using the ‘haplotype’ function of the package ‘haplotypes’ (Aktas, 2015). We calculated haplotype diversity of each population (Nei and Tajima, 1981) with the function ‘hap.div’, and nucleotide diversity (Nei, 1987) with the function ‘nuc.div’, both available in the ‘pegas’ package (Paradis, 2010). We tested for significant differences in the genetic diversity of the North and Baltic Sea by conducting Analyses of Variance (ANOVA) for each species and diversity measure using custom scripts available online (10.6084/m9.figshare.c.5341910). We calculated Tajima’s D and its deviation from zero with the function ‘tajima.test’ of the ‘pegas’ package (Paradis, 2010). Tajima’s D is the test statistic that calculates the difference between the expected genetic diversity based on the number of segregating sites and the average number of pairwise differences (Tajima, 1989). A negative Tajima’s D indicates either a recent selective sweep or population expansion after a bottleneck, the expectation for a relatively recent colonization.

We calculated genetic differentiation between North and Baltic Sea populations as Φ_ST_ with the function ‘pairwiseTest’ of the package ‘strataG’ (Archer et al., 2017), Jost D with the function ‘pairwise_D’ of the ‘mmod’ package (Winter, 2012) and wrote our own function to calculate the nearest neighbor statistic Snn (Hudson, 2000). Φ_ST_ is a derivative of the classical fixation index F_ST_, adapted for mitochondrial haplotype data (Excoffier et al., 1992). Jost D is supposed to be a more accurate measure of population differentiation when genetic diversity is high and the number of unique alleles per population is large (Jost, 2008). Snn is particularly powerful when sample sizes are small or uneven between populations (Hudson, 2000). We estimated significant deviations from zero (no differentiation between population pairs) for all differentiation indices by comparing the point estimates with an empirical distribution of values based on 1000 permutations.

### Rarefaction analysis

Initially, we included species for which at least five sequences for each population were available. This low number is sufficient to distinguish high- and low-diversity populations (Goodall-Copestake et al., 2012). To ensure that any observed lack of genetic differentiation was not due to small sample size (i.e. lack of power), we randomly subsampled all species in which populations had more than 20 sequences to 5, 10 or 15 sequences per population, and re-calculated population genetic estimates on these random subsamples. We repeated the subsampling 100 times, and compared the distribution of these estimates with the point estimates of the full dataset for each species. We also repeated all population genetic analyses on datasets rarefied to the same number of sequences per population. In this last iteration, different species can have different sample sizes, but the sample sizes are the same for each population within a species.

### Coalescent estimates of theta and migration rate

Differentiation indices such as Φ_ST_ assume, for example, that migration rates are symmetric. The coalescent approach, on the other hand, allows migration rates to vary (Beerli, 2006), which is relevant for testing the hypothesis of the Baltic Sea as a population sink. For all species with more than 20 sequences per population, we estimated the mutation-rate scaled migration rates between North and Baltic Sea populations m, the mutation-rate scaled effective population size q of each population and the time since divergence t, implemented in the software ‘IMa2’ v.8.27.12 (Hey and Nielsen, 2004). IMa2 uses Bayesian inference to estimate posterior probability densities of these population genetic parameters. It is particularly well-suited for populations that diverged recently (Hey and Nielsen, 2004). We used the HKA model of sequence evolution, uniform priors, and the low heating scheme described in the IMa2 manual, and replicated each run five times to confirm convergence. For details on each species, see the output files that are available online (10.6084/m9.figshare.c.5341910). Convergence was further ensured by high effective sampling sizes (ESS), and single posterior probability density peaks. Divergence times were converted to years by dividing the divergence estimates by the mutation rate per year scaled to the respective alignment length. We based this mutation rate on a substitution rate of 1.22% per one million years, which appears to be similar across marine invertebrates (Wilke et al., 2009).

### Correlations between genetic and life history data

Given the very short time that non-native species have been present in the Baltic Sea, we do not expect them to follow the same rules as native species, who evolved in the North Sea - Baltic Sea system. We therefore excluded non-native species from the following analyses. We estimated the effects of life history on either genetic differentiation between North and Baltic Sea or on salinity tolerance, which we approximated by the easternmost longitude a species was recorded from in the Baltic Sea, as well as the effects of these two diversification measurements to each other. We used the Bayesian approach implemented in the ‘MCMCglmm’ package using the ‘mcmcglmm’ function (Hadfield, 2010). All life history traits were treated as fixed effects to assess their significance. In particular, we included the dispersal potential of larvae, the dispersal potential of adults, and the taxon into the models. Given that these models are overparameterized for our sample size, the number of species, we sequentially removed response variables that were non-significant from the model. We used the default priors, and let the model run for 60,000 generations. Significance was assessed using the mcmc p-value (pMCMC). To ensure convergence, we inspected the traces and checked the posterior densities visually for normality, and made sure the effective sample size was larger than 200 in all variables.

## Results

### Acquisition of life history data

Life history data were retrieved from studies published throughout the 20^th^ and 21^st^ centuries, the earliest ones dating back more than 100 years. These historical studies may not always meet modern methodological standards, but were for several species the only available source of information. For little studied species and/or species that are difficult to rear/brood in the laboratory (i.e. *Balanus crenatus, Ophiura albida, Palaemon varians*), we had to estimate PLD and salinity tolerance from studies not particularly addressing these traits. We decided to generally report salinity tolerance data for adult organisms (Tab. 1), even though the larvae of many marine invertebrates are known to require higher salinities to undergo full development (see e.g. Anger, 2001 for crustaceans; Sherman et al., 2016). However, larval salinity tolerances were available for very few of the investigated species only, and were lacking for Baltic Sea populations (with the exception of *C. maenas*, see below).

**Table 1.**
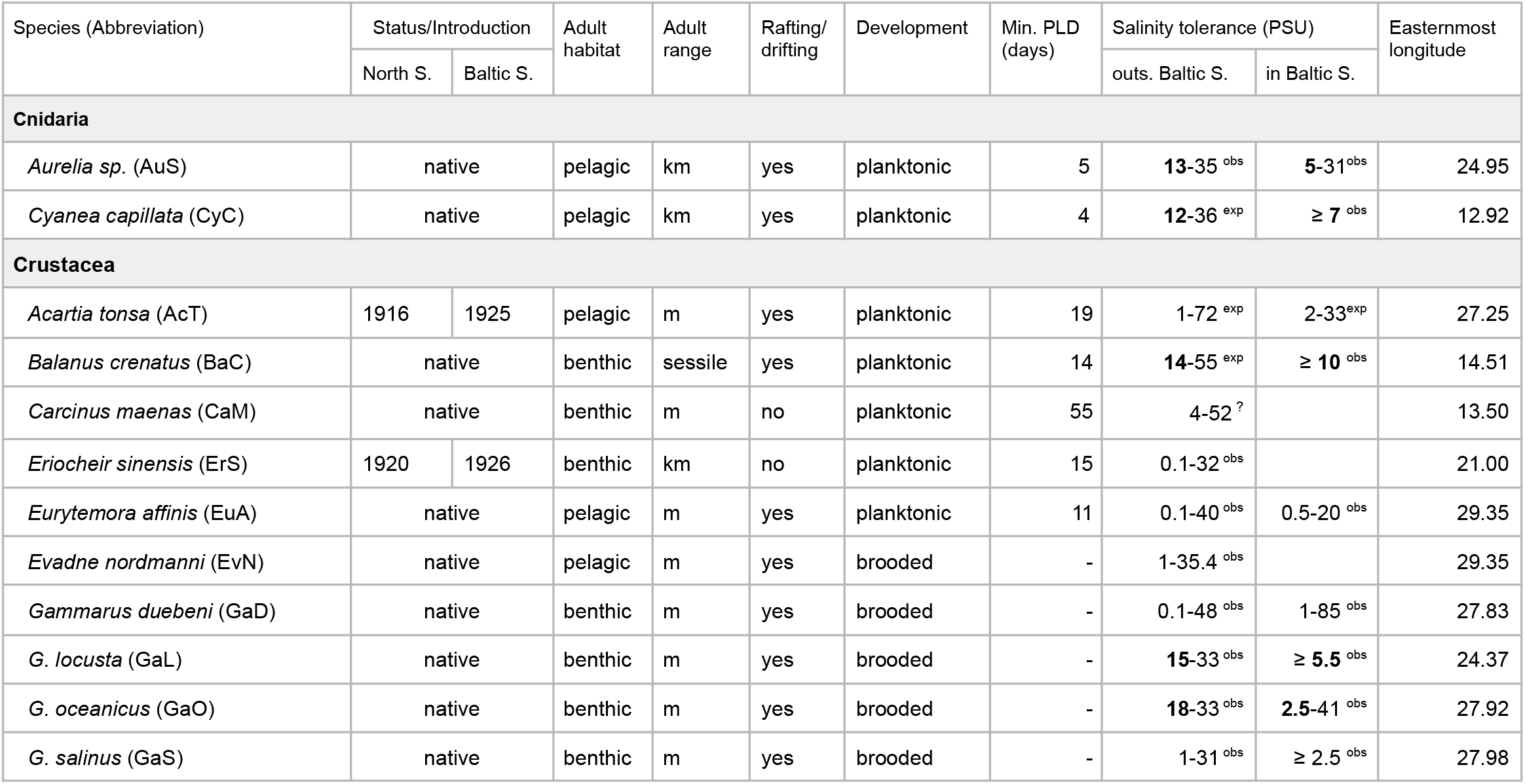
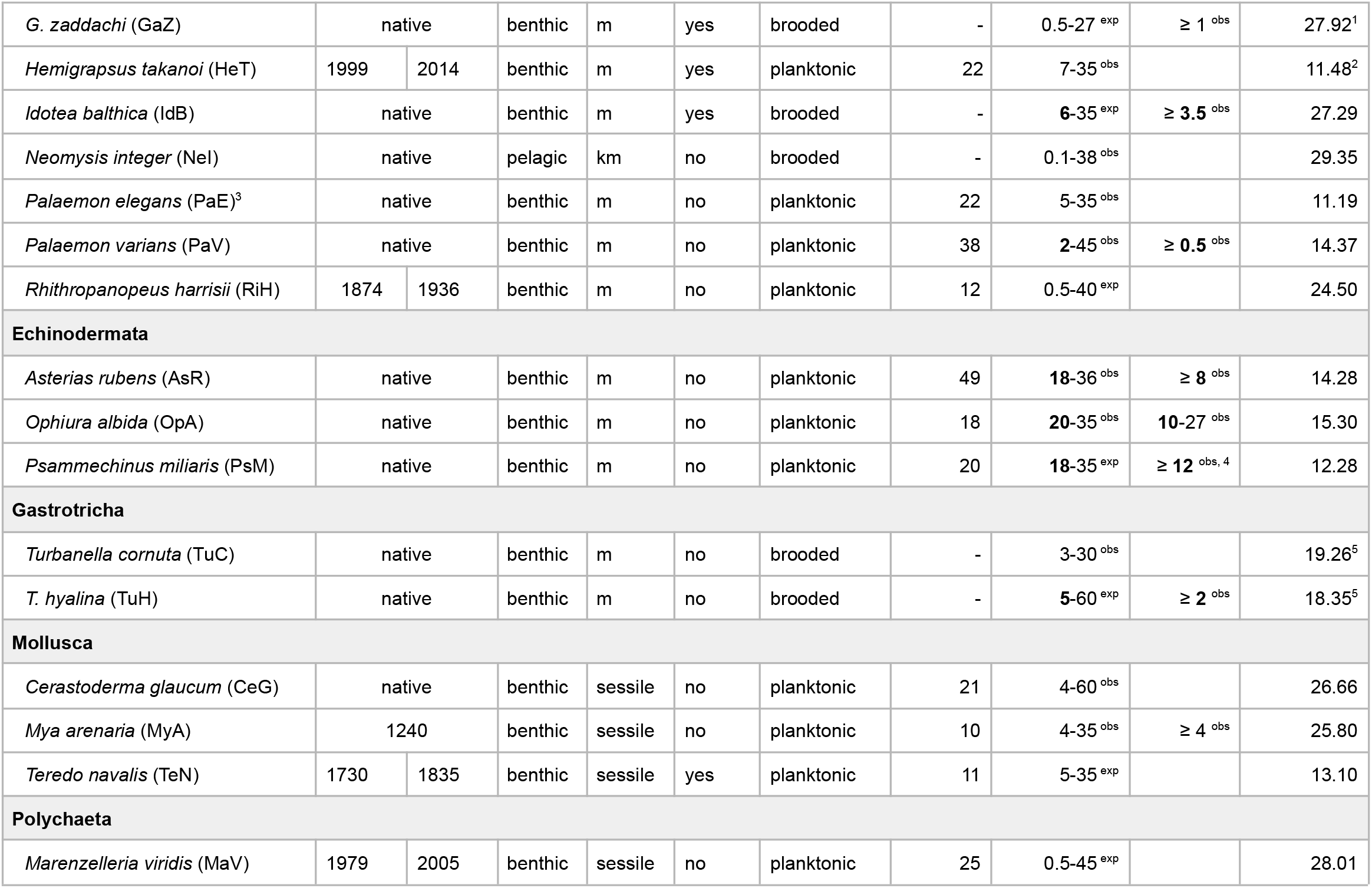
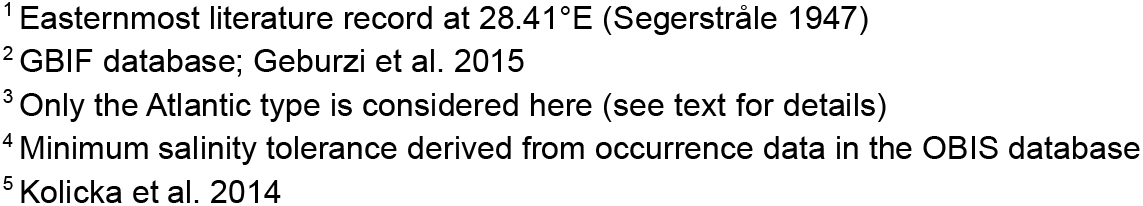
Life history traits of the investigated species. For non-native species, the year of the first record for North and Baltic Seas is given. Salinity tolerances are derived from either experimental (exp) or observational (obs) studies. They are given for populations outside the Baltic Sea and, where available, Baltic Sea populations; bold numbers highlight differences in lower salinity tolerance limits in- and outside the Baltic Sea of ≥ 1 PSU. Easternmost longitudes are derived from the OBIS database except for three species (see footnotes).

| Species (Abbreviation) | Status/Introduction |  | Adult habitat | Adult range | Rafting/ drifting | Development | Min. PLD (days) | Salinity tolerance (PSU) |  | Easternmost longitude |
| --- | --- | --- | --- | --- | --- | --- | --- | --- | --- | --- |
|  | North S. | Baltic S. |  |  |  |  |  | outs. Baltic S. | in Baltic S. |  |
| Cnidaria |  |  |  |  |  |  |  |  |  |  |
| *Aurelia sp.* (AuS) | native |  | pelagic | km | yes | planktonic | 5 | **13-35**<sup>obs</sup> | **5-31**<sup>obs</sup> | 24.95 |
| *Cyanea capillata* (CyC) | native |  | pelagic | km | yes | planktonic | 4 | **12-36**<sup>exp</sup> | **≥ 7**<sup>obs</sup> | 12.92 |
| Crustacea |  |  |  |  |  |  |  |  |  |  |
| *Acartia tonsa* (AcT) | 1916 | 1925 | pelagic | m | yes | planktonic | 19 | 1-72<sup>exp</sup> | 2-33<sup>exp</sup> | 27.25 |
| *Balanus crenatus* (BaC) | native |  | benthic | sessile | yes | planktonic | 14 | **14-55**<sup>exp</sup> | **≥ 10**<sup>obs</sup> | 14.51 |
| *Carcinus maenas* (CaM) | native |  | benthic | m | no | planktonic | 55 | 4-52<sup>?</sup> |  | 13.50 |
| *Eriocheir sinensis* (ErS) | 1920 | 1926 | benthic | km | no | planktonic | 15 | 0.1-32<sup>obs</sup> |  | 21.00 |
| *Eurytemora affinis* (EuA) | native |  | pelagic | m | yes | planktonic | 11 | 0.1-40<sup>obs</sup> | 0.5-20<sup>obs</sup> | 29.35 |
| *Evadne nordmanni* (EvN) | native |  | pelagic | m | yes | brooded | - | 1-35.4<sup>obs</sup> |  | 29.35 |
| *Gammarus duebeni* (GaD) | native |  | benthic | m | yes | brooded | - | 0.1-48<sup>obs</sup> | 1-85<sup>obs</sup> | 27.83 |
| *G. locusta* (GaL) | native |  | benthic | m | yes | brooded | - | **15-33**<sup>obs</sup> | **≥ 5.5**<sup>obs</sup> | 24.37 |
| *G. oceanicus* (GaO) | native |  | benthic | m | yes | brooded | - | **18-33**<sup>obs</sup> | **2.5-41**<sup>obs</sup> | 27.92 |
| *G. salinus* (GaS) | native |  | benthic | m | yes | brooded | - | 1-31<sup>obs</sup> | **≥ 2.5**<sup>obs</sup> | 27.98 |
| *G. zaddachi* (GaZ) | native |  | benthic | m | yes | brooded | - | 0.5-27 <sup>exp</sup> | ≥ 1 <sup>obs</sup> | 27.92<sup>1</sup> |
| *Hemigrapsus takanoi* (HeT) | 1999 | 2014 | benthic | m | yes | planktonic | 22 | 7-35 <sup>obs</sup> |  | 11.48<sup>2</sup> |
| *Idotea balthica* (IdB) | native |  | benthic | m | yes | brooded | - | 6-35 <sup>exp</sup> | ≥ 3.5 <sup>obs</sup> | 27.29 |
| *Neomysis integer* (NeI) | native |  | pelagic | km | no | brooded | - | 0.1-38 <sup>obs</sup> |  | 29.35 |
| *Palaemon elegans* (PaE)<sup>3</sup> | native |  | benthic | m | no | planktonic | 22 | 5-35 <sup>obs</sup> |  | 11.19 |
| *Palaemon varians* (PaV) | native |  | benthic | m | no | planktonic | 38 | 2-45 <sup>obs</sup> | ≥ 0.5 <sup>obs</sup> | 14.37 |
| *Rhithropanopeus harrisii* (RiH) | 1874 | 1936 | benthic | m | no | planktonic | 12 | 0.5-40 <sup>exp</sup> |  | 24.50 |
| Echinodermata |  |  |  |  |  |  |  |  |  |  |
| *Asterias rubens* (AsR) | native |  | benthic | m | no | planktonic | 49 | 18-36 <sup>obs</sup> | ≥ 8 <sup>obs</sup> | 14.28 |
| *Ophiura albida* (OpA) | native |  | benthic | m | no | planktonic | 18 | 20-35 <sup>obs</sup> | 10-27 <sup>obs</sup> | 15.30 |
| *Psammechinus miliaris* (PsM) | native |  | benthic | m | no | planktonic | 20 | 18-35 <sup>exp</sup> | ≥ 12 <sup>obs, 4</sup> | 12.28 |
| Gastrotricha |  |  |  |  |  |  |  |  |  |  |
| *Turbanella cornuta* (TuC) | native |  | benthic | m | no | brooded | - | 3-30 <sup>obs</sup> |  | 19.26<sup>5</sup> |
| *T. hyalina* (TuH) | native |  | benthic | m | no | brooded | - | 5-60 <sup>exp</sup> | ≥ 2 <sup>obs</sup> | 18.35<sup>5</sup> |
| Mollusca |  |  |  |  |  |  |  |  |  |  |
| *Cerastoderma glaucum* (CeG) | native |  | benthic | sessile | no | planktonic | 21 | 4-60 <sup>obs</sup> |  | 26.66 |
| *Mya arenaria* (MyA) | 1240 |  | benthic | sessile | no | planktonic | 10 | 4-35 <sup>obs</sup> | ≥ 4 <sup>obs</sup> | 25.80 |
| *Teredo navalis* (TeN) | 1730 | 1835 | benthic | sessile | yes | planktonic | 11 | 5-35 <sup>exp</sup> |  | 13.10 |
| Polychaeta |  |  |  |  |  |  |  |  |  |  |
| *Marenzelleria viridis* (MaV) | 1979 | 2005 | benthic | sessile | no | planktonic | 25 | 0.5-45 <sup>exp</sup> |  | 28.01 |

<sup>1</sup> Easternmost literature record at 28.41°E (Seggerstråle 1947)
<sup>2</sup> GBIF database; Geburzi et al. 2015
<sup>3</sup> Only the Atlantic type is considered here (see text for details)
<sup>4</sup> Minimum salinity tolerance derived from occurrence data in the OBIS database
<sup>5</sup> Kolicka et al. 2014

When retrieving data of the species’ eastern distribution limits in the Baltic Sea from OBIS or GBIF, we excluded isolated data points (e.g. for *Psammechinus miliaris*), as well as data points prior to 1990 (e.g. for *Asterias rubens*). These old data points are excluded because salinity may have changed, and we want a comparable picture of salinity tolerance. The shrimp *Palaemon elegans* is a special case, as two genetically highly divergent lines occur in the Baltic Sea. One of them is also present in the North Sea and Atlantic (the Atlantic type), and one was most likely introduced to the Baltic Sea from the Black Sea (the Black Sea type) (Reuschel et al., 2010). It is unclear if these lineages hybridize. Their genetic distance suggests that they are separate species. We only consider the Atlantic lineage here, as the non-native Black Sea lineage does not (yet) occur in the North Sea (pers. comm. A. Böttcher). We therefore disregarded life history and distribution data from regions in the Baltic Sea where the Black Sea type occurs according to Reuschel et al. (2010).

The majority of species (22) is benthic, their adult mobility typically being in the range of meters. Five species (*Balanus crenatus, Cerastoderma glaucum, Marenzelleria viridis, Mya arenaria*, and *Teredo navalis*) are sessile as adults. A planktonic larval phase occurs in 18 of the investigated species, four of them being pelagic and 14 being benthic as adults, including all sessile species. Pelagic larval duration (PLD) ranges from less than a week in the cnidarians *Aurelia sp*. and *C. capillata* to six weeks and more in *A. rubens* and *C. maenas* (Tab. 1). The investigated species thus cover a broad range of dispersal potentials when jointly considering adult mobility and the presence and duration of a planktonic larval phase.

At the upper extreme are the two cnidarian species, with highly mobile, pelagic adults and planktonic larvae. Among the benthic species, *E. sinensis* stands out with highly mobile adults, capable of long-distance migrations, and a PLD of about two weeks. Ten further benthic species have a PLD exceeding two weeks. In all *Gammarus* species, as well as *Idotea balthica*, long-range dispersal of adults may occur by individuals rafting on floating macroalgae or seagrass, potentially ‘boosting’ adult mobility under favorable conditions. At the lower extreme are the two gastrotrich species that live in-benthic as adults, attach their eggs to sand grains within the sediment, and lack a planktonic larval phase.

Most of the investigated species have broad to very broad salinity tolerances, ranging from (nearly) freshwater to fully marine or even hypersaline conditions. In general, the further east a species occurs into the Baltic Sea (i.e. the more brackish conditions it tolerates), the broader its salinity range appears to be (Fig. 2). For *Aurelia sp*., *Cyanea capillata, Balanus crenatus, Gammarus locusta, G. oceanicus, Idotea balthica, P. varians, Asterias rubens, Ophiura albida, Psammechinus miliaris* and *Turbanella hyalina*, we found explicit literature evidence that their Baltic Sea populations tolerate lower salinities compared to North Sea/Atlantic populations (Tab. 1). This difference was most pronounced in *G. oceanicus, A. rubens* and *O. albida*, with their Baltic Sea populations tolerating water less saline by 10 PSU and more compared to their North Sea populations. *Carcinus maenas* is a special case when comparing salinity tolerances, as adults from North and Baltic Sea populations do not seem to differ in their salinity tolerances, but the Baltic Sea populations of *C. maenas* are able to complete larval development at much lower salinities compared to their North Sea conspecifics (13 vs. 20 PSU; Dries and Adelung, 1982; Anger et al., 1998).

**Figure 2.**
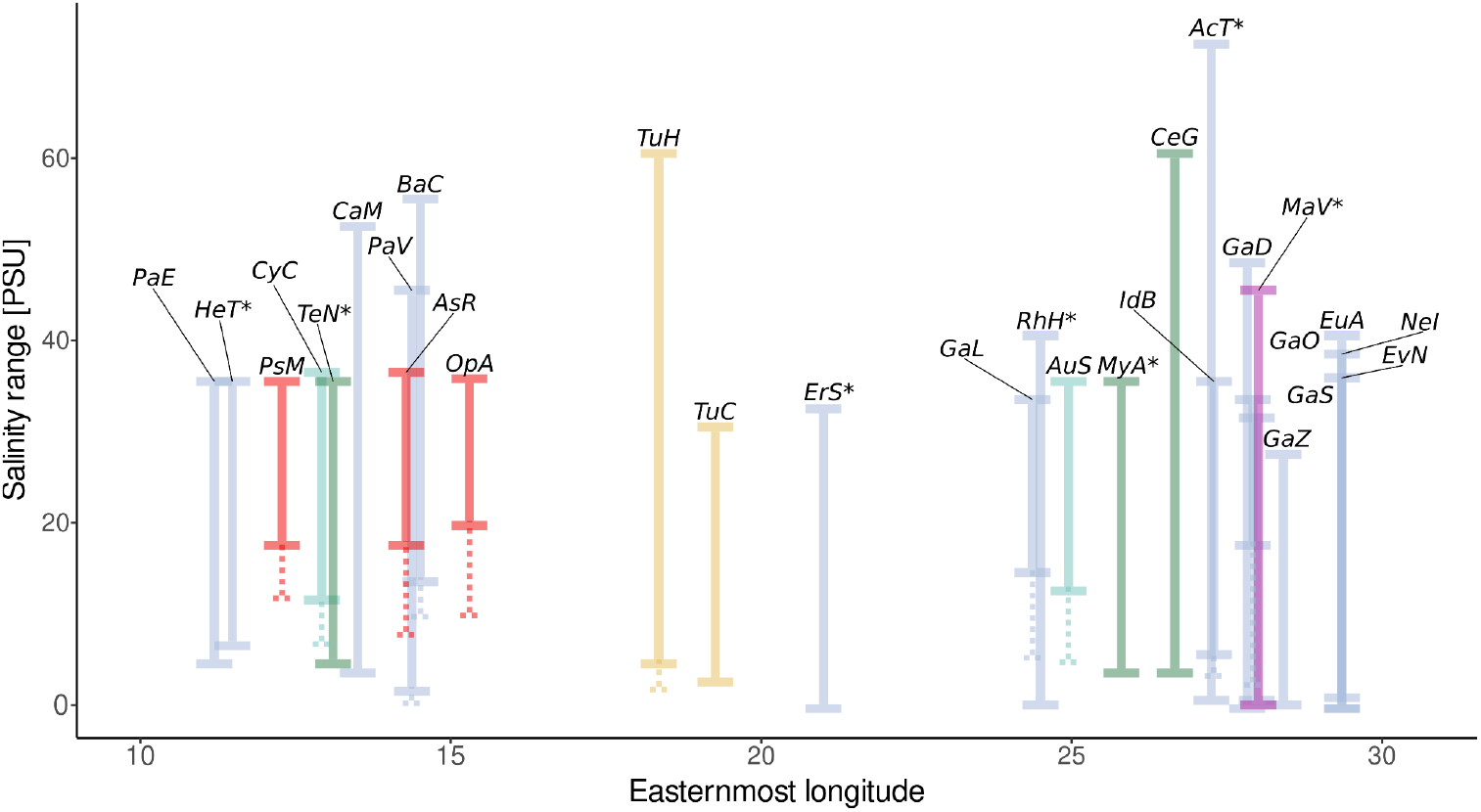
Salinity tolerances and eastern distribution limit in the Baltic Sea of the investigated species. Dotted lines indicate enhanced low-salinity tolerance in Baltic Sea populations in species where Baltic Sea-specific salinity tolerance data were available. For species abbreviations, see Tab 1.

Seven of the investigated species are considered non-native in the Baltic Sea: *Acartia tonsa, Eriocheir sinensis, Hemigrapsus takanoi, Rhithropanopeus harrisii, Mya arenaria, Teredo navalis* and *Marenzelleria viridis*. Most non-native species have a long planktonic larval phase (PLD > 2 weeks, except *R. harrisii* and *M. arenaria*; Tab. 1), a typical trait of successful invasive marine invertebrates. Time since introduction in the Baltic Sea varies between almost 800 years (*Mya arenaria*; Petersen et al., 1992; Behrends et al., 2005) and six years (*Hemigrapsus takanoi*; Geburzi et al., 2015). The easternmost longitude of regular occurrence in the Baltic Sea corresponds fairly well with the salinity tolerance for most of the non-native species. Only the recently established *H. takanoi* has its current eastern distribution limit distinctly west of the 7 PSU isohaline (compare Fig. 1 and Fig. 5).

### Acquisition of genetic data

Since the review of Johannesson and André (2006) on Baltic Sea genetic diversity, much new COI data have been generated (Tab. S2). Sequences for North Sea populations were abundant, and generally not the limiting factor. A good source for North Sea data were recent large-scale barcoding efforts for Crustacea (Raupach et al., 2015), Mollusca (Barco et al., 2016) and Echinodermata (Laakmann et al., 2016). We excluded data from several studies that sequenced different mitochondrial fragments, or that did not provide enough information to reconstruct haplotype frequencies (e.g. *Gammarus tigrinus*: Kelly et al., 2006; *Limecola balthica*: Becquet et al., 2012). In the case of the shipworm *Teredo navalis*, we included three locations that were sampled after 2012, and were not close to each other: Kiel, Kühlungsborn and Hiddensee, to thin the otherwise very large number of sequences. In accordance with the life history data, we only use COI data from the Atlantic lineage of *Palaemon elegans* (see above). We generated new data for the Baltic Sea populations of seven species: three echinoderms, three brachyuran crabs, one caridean shrimp and one barnacle (Tab. S2). By combining sequence data from online resources with this newly generated data, we were able to consider 28 species, including members of the Cnidaria (2 spp.) Crustacea (17 spp.), Echinodermata (3 spp.), Gastrotricha (2 spp.), Mollusca (3 spp.) and Polychaeta (1 sp.) (Fig. 3, Tab. S2). In most cases, the barcoding marker located at the 5’ end of the COI gene was amplified (Folmer et al., 1994). Population sample sizes ranged from six to 196 sequences with a median of 25 sequences, and alignment lengths varied from 423 to 675 bp with a mean of 545 bp. For eight species, sequences were available for the transition zone as defined in Fig. 1. Of these species, three were highly differentiated: *Cerastoderma glaucum, Eurytemora affinis* and *Balanus crenatus*. This allowed us to clearly assign the transition zone sequences to either the North or Baltic Sea population (see Fig. S3.1 in the Supporting Information 3). From this, we inferred the break between North and Baltic Sea populations: between the Limfjord and North Sea (*C. glaucum*), between Skagerrak and Kattegat (*B. crenatus*) and between Skagerrak and the North Sea (*E. affinis*). In the case of *E. affinis*, we also included the few available White Sea sequences, which clustered with the North Sea sequences.

**Figure 3.**
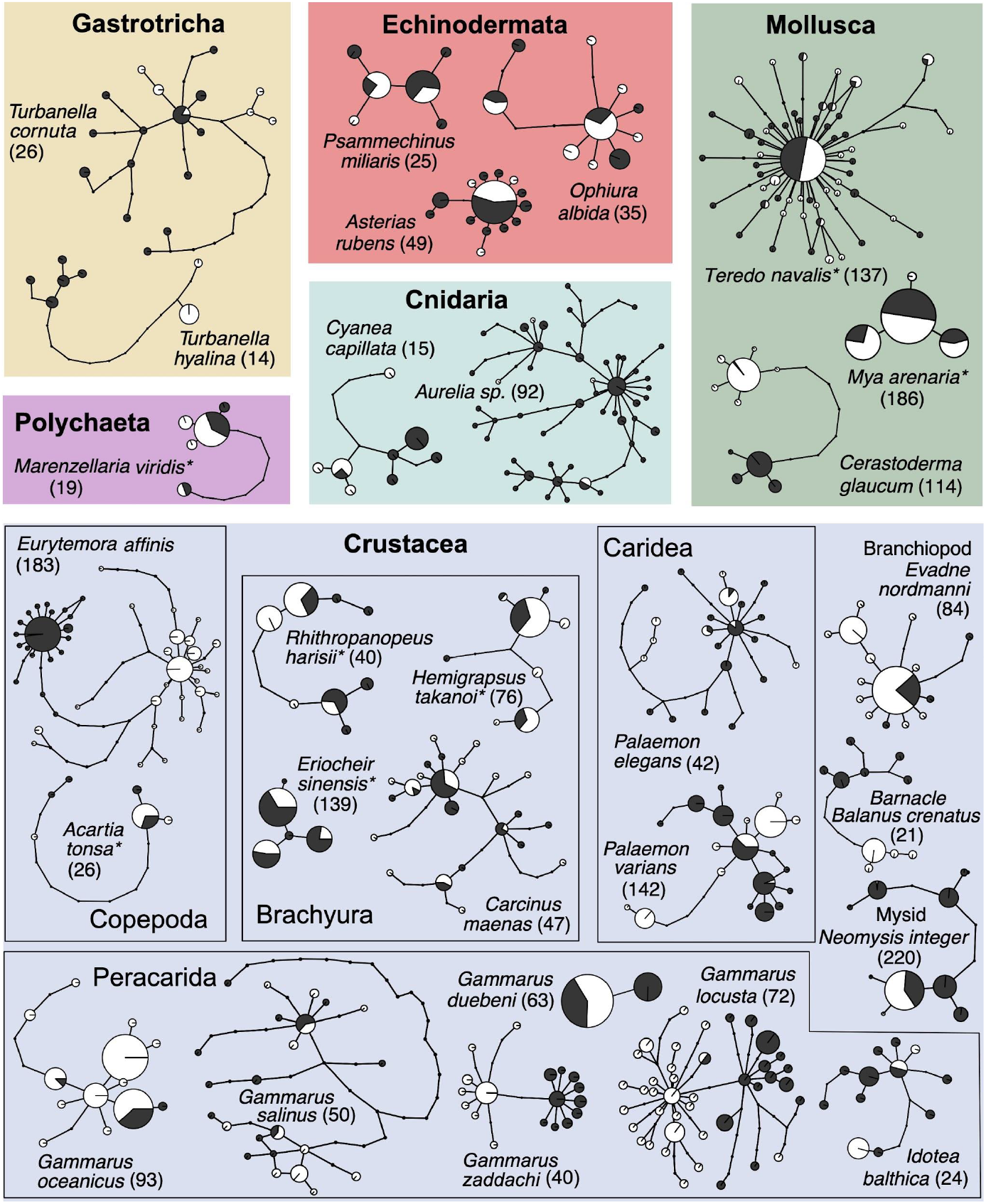
Haplotype networks for all 28 investigated species grouped by taxonomic affinity. Size of the circles is relative to the sample size, but not the same between species. Colors denote populations: black = North Sea, white = Baltic Sea, and asterisks denote non-native species. Overall sample size n in parentheses.

### Rarefaction analysis

Calculating genetic differentiation from only 5 sequences per population for 10 species with more than 20 available sequences per population generated large ranges of differentiation index values. For Φ_ST_, the 95% range of estimated values was 0.83 averaged across all 10 species, and for Snn, this interval was 0.36 (see Fig. S3.2 in the Supporting Information 3). Given that Φ_ST_ ranges from 0 to 1 and Snn from 0.5 to 1, these ranges are comparably wide. While the range was wide, the majority of estimates centered around the value calculated from the full data set. In some species, we observed an upward bias, such that Φ_ST_ and Snn would be larger when sample sizes are small. The range for species with very high genetic differentiation (*Cerastoderma glaucum, Eurytemora affinis* and *Gammarus locusta*) was much lower, suggesting that for highly differentiated species, small sample sizes provide accurate results. Increasing the sample size reduced the 95% intervals of estimated Φ_ST_ and Snn values (see Fig. S3.2), but variability of Φ_ST_ values reduced more with larger sample sizes than variability of Snn values.

Measures of genetic diversity were relatively robust to small sample size with relatively small ranges (Fig. S3.2). Nucleotide diversity appeared to be a robust measure of genetic diversity. In contrast, Tajima’s D had a wide range of values even at larger sample sizes (Fig. S3.2). In some instances, the ranges did not even include the value estimated from the full dataset. Thus this test statistic is only useful when sample sizes are large in comparison to the number of haplotypes. Considering that Tajima’s D is based on the abundance of rare alleles, this result is expected. As a result of these rarefaction analyses, Φ_ST_ will be considered as the most robust measure of differentiation in the subsequent analyses, and nucleotide diversity the most robust point estimate of diversity. Tajima’s D, on the other hand, will be considered less valuable for species with small sample sizes. We did not remove such species as they may add to our understanding of the role of the Baltic Sea as a diversification hotspot, especially when they are strongly differentiated.

### Population genetic analyses

Genetic diversity varied considerably between species and populations. Only five species had even haplotype diversities in the North and Baltic Sea, whereas eight species had higher haplotype diversity in the Baltic Sea, and 13 species had lower haplotype diversity in the Baltic Sea (Fig. 4A). The Baltic Sea population of the amphipod *Gammarus duebeni* had a haplotype diversity of 0, which means only a single haplotype was found in the Baltic Sea. This species also had only two haplotypes in the North Sea, making its genetic diversity extremely low (compare Fig. 3). No North Sea population, on the other hand, had a haplotype diversity lower than 0.4. With regards to nucleotide diversity, 14 species were equally diverse in the North and Baltic Sea, and five species were more diverse in the Baltic Sea than the North Sea (Fig. 4B). The remaining eight species were more diverse in the North Sea. The ratio of nucleotide diversities from North and Baltic Sea populations was strongly correlated with the respective ratio of haplotype diversities (Pearson’s product-moment correlation = 0.722, 95% confidence interval = 0.472, 0.865, p-value < 0.001). Only the amphipod *Gammarus locusta* had a significantly higher haplotype diversity in the Baltic Sea than in the North Sea, but the nucleotide diversity showed the opposite trend.

**Figure 4.**
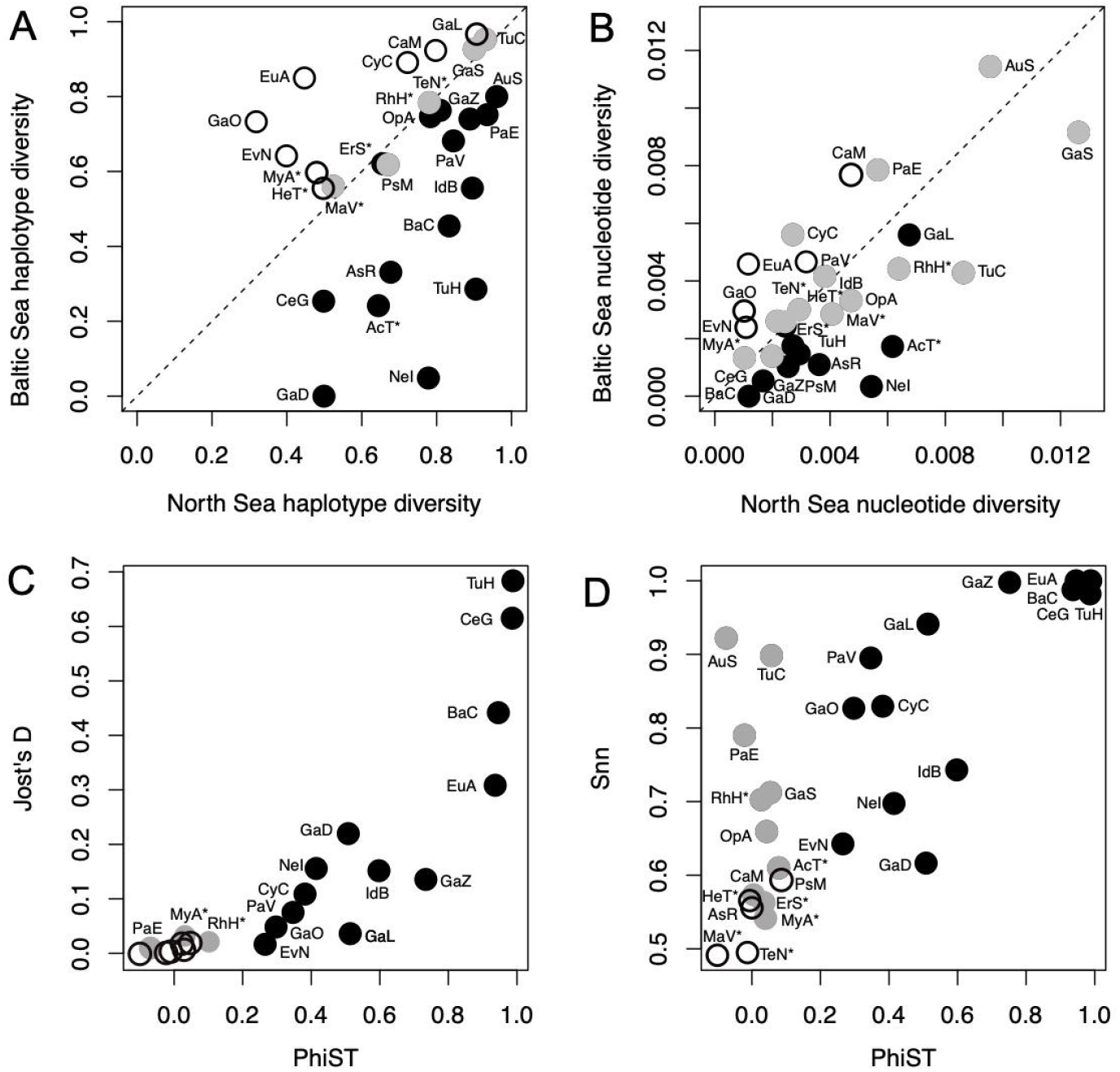
Population genetic comparison of North and Baltic Sea populations of marine invertebrates. A: Haplotype diversity, B: nucleotide diversity. Black: more diverse in North Sea, white: more diverse in Baltic Sea, grey: equally diverse. C: Comparison between the differentiation indices Φ_ST_ and Jost’s D, D: comparison between the differentiation indices Φ_ST_ and Hudson’s Snn. Black: significantly differentiated with both indices, grey: only differentiated with Jost’s D (C) or Hudson’s Snn (D), white: undifferentiated with either index. For species abbreviations see Tab 1.

The two differentiation indices Φ_ST_ and Jost’s D gave qualitatively similar results (Fig. 4C). Φ_ST_ identified about half of the species as significantly differentiated. Jost’s D identified three additional species, the shrimp *Palaemon elegans*, and the two non-native species with the oldest introduction date, the soft shell clam *Mya arenaria* and the crab *Rhithropanopeus harrisii*, as significantly differentiated. Hudson’s Snn identified five additional species as significantly differentiated for a total of 22 species (Fig. 4D). Five species were considered undifferentiated by all test statistics, three non-native species (Fig. 4C, D), and two native echinoderms, the sea star *A. rubens* and the sea urchin *Psammechinus miliaris*.

For 17 species, Tajima’s D was not significantly different from zero for either North Sea or Baltic Sea population (see Fig. S3.3 in the Supporting Information 3). Four species (*Gammarus locusta, Turbanella hyalina, Acartia tonsa, Neomysis integer*) had significantly negative values for the Baltic Sea population, while *Palaemon varians* had a significantly positive value in the Baltic Sea. *Evadne nordmanni* had a significantly negative value in the North Sea. Two species had Tajima’s D values that were smaller than zero in both populations: *Teredo navalis* and *Eurytemora affinis*. The large number of insignificant values matches our rarefaction analysis, which showed that only large sample sizes estimate Tajima’s D with confidence.

### Coalescent estimates of theta and migration rate

More than 20 sequences per population were available for 12 of the 28 species (Tab. S2). Of those, four were non-native species, for which we could neither obtain high effective sample sizes (ESS) nor reliable estimates of population size and migration rates (supplementary figures S4 & S5). We are violating the coalescent assumptions grossly in those populations: they did not evolve their current genetic diversity in the North and Baltic Sea. In contrast, a presumably random subset of variants was translocated to the non-native range, with possible mixing of genetically divergent populations. Thus we cannot put any confidence into the obtained population size and migration rate estimates for those four species. Two other species, the cladoceran *Evadne nordmanni* and the shore crab *Carcinus maenas*, had ESS lower than 200 for at least some estimates, and marginal posterior probability distribution with multiple peaks.

The remaining six species converged on specific estimates with single peaks in the marginal posterior probability and ESS larger than 1300 for all estimates (Fig. S3.4, S3.5). For the cockle *Cerastoderma glaucum*, the copepod *Eurytemora affinis* and the amphipod *Gammarus locusta*, population sizes were larger in the Baltic Sea than in the North Sea, whereas the opposite was the case for the amphipod *Gammarus duebeni*,the shrimp *Palaemon varians* and the opossum shrimp *Neomysis integer* (Fig. S4). Migration rates were larger from the Baltic Sea to the North Sea in the cockle *C. glaucum*, the amphipod *G. locusta* and the opossum shrimp *N. integer*, and larger from the North Sea to the Baltic Sea for the copepod *E. affinis* and the shrimp *P. varians*. The migration rates for the amphipod *G. duebeni* were very wide (Fig. S5), which is not surprising given the extremely low genetic diversity with a single segregating site.

IMa2 also estimated divergence times between populations. The amphipod *G. duebeni* had the most recent estimate with 1524 years, and a 95% highest posterior density (HPD) interval from 197 to 1593 years. The other four species had wide 95% HPD intervals: 98,004–1,570,498 (*Palaemon varians*), 201,192–450,820 years (*G. locusta*), 232,456–1,492,743 years (*E. affinis*), and 146,601–1,581,838 years (*C. glaucum*) when assuming a generation time of one year for all species but the copepod *E. affinis*, for which we assumed a generation of ½ year. These confidence intervals overlap between 200,000 and 450,000 years ago, indicating a possible concurrent colonization event at that time.

### Correlations between genetic differentiation, salinity tolerance and life history data

Life history and dispersal traits did not have significant linear relationships with the level of genetic differentiation. The MCMCglmm results showed that Φ_ST_ itself was significantly larger than zero across all native species, indicating significant differentiation across the North and Baltic Sea (pMCMC = 0.000335). Conversely, higher taxonomic units, adult mobility or minimum PLD did not significantly correlate with Φ_ST_ (Fig. 5A). However, the species with the longest PLD were little differentiated (*Carcinus maenas, Asterias rubens*) (Fig. 5A). Genetic differentiation varied greatly between species without planktonic larvae. For species with planktonic larvae, genetic differentiation was either absent or very pronounced (Fig. 5A). It does stand out that none of the investigated Echinodermata and almost none of the Mollusca were significantly differentiated between the North and Baltic Sea. All investigated alien species have low levels of population differentiation (Fig. 4), which is expected given their relatively recent introduction into the Baltic Sea.

**Figure 5.**
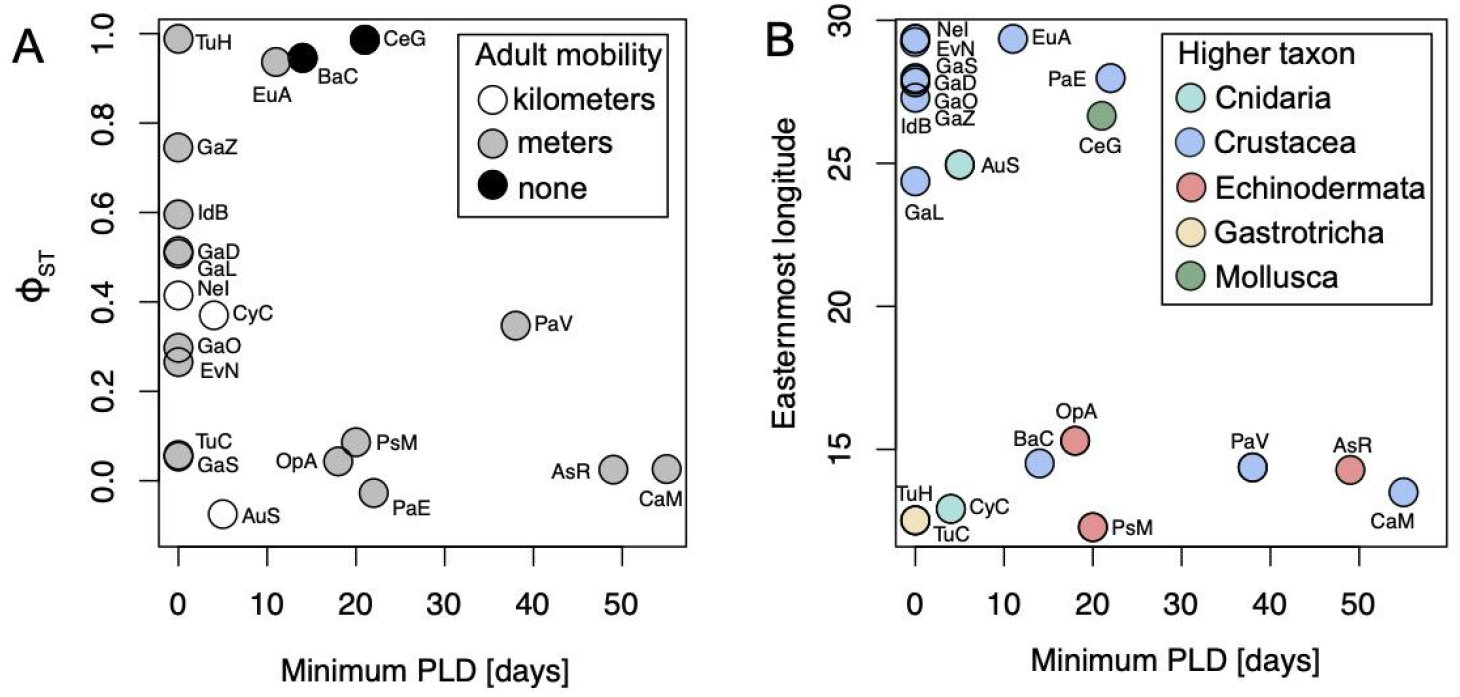
Life history and genetic differentiation. Genetic differentiation vs PLD, and genetic differentiation vs easternmost longitude (which correlates strongly with salinity, see Fig. 1).

The easternmost longitude at which a species was reported in the Baltic Sea is a proxy of its natural salinity tolerance, as salinity declines to the east (Fig. 1). This easternmost longitude was significantly and negatively affected by the minimum PLD (pMCMC = 0.01122), and the taxonomic affinity (Fig. 5B). In particular, Mollusca were found further to the east (pMCMC = 0.00536), although this result is based on a single species, the cockle *Cerastoderma glaucum*.

## Discussion

In this comparative study, we compiled genetic, ecological and life history data for 28 marine invertebrate species that occur in both North and Baltic Sea. These species live under the marine conditions of the North Sea, and under the brackish conditions of the Baltic Sea. We asked whether these species have broad ecological tolerances, and are swept into the Baltic Sea without adapting or diversifying, or whether the Baltic Sea acts as a diversification hotspot with low gene flow to the North Sea. Taking all of the available evidence together, we identified significant ecological and/or genetic differentiation for 18 of the 28 investigated species (Fig. 6). For these 18 species, the Baltic Sea represents a diversification hotspot. The Baltic Sea appears to be a population sink for ten species, seven of which are non-natives (Fig. 6). For these seven non-natives, the lack of population differentiation and often equal genetic diversity between North and Baltic Sea are the result of their recent expansion into the Baltic Sea rather than long term population sink dynamics. Moreover, for the majority of non-native species, no data on ecological diversification exists. The native gastrotrich *Turbanella cornuta*, which does not show signs of genetic differentiation, has also not been assessed for ecological diversification. The shrimp *Palaemon elegans* is significantly differentiated using two of the three differentiation indices, and the amphipod *Gammarus salinus* is differentiated based on Hudson’s Snn. Thus for 85% of the native species, and potentially all of them, the Baltic Sea represents a diversification hotspot, irrespective of their dispersal potential. It may well be that all species which successfully colonized the Baltic Sea had to diversify. Investigating the non-native species may provide clues as to the timing of such events, as would probing the genomes of natives for signatures of adaptive evolution.

**Figure 6.**
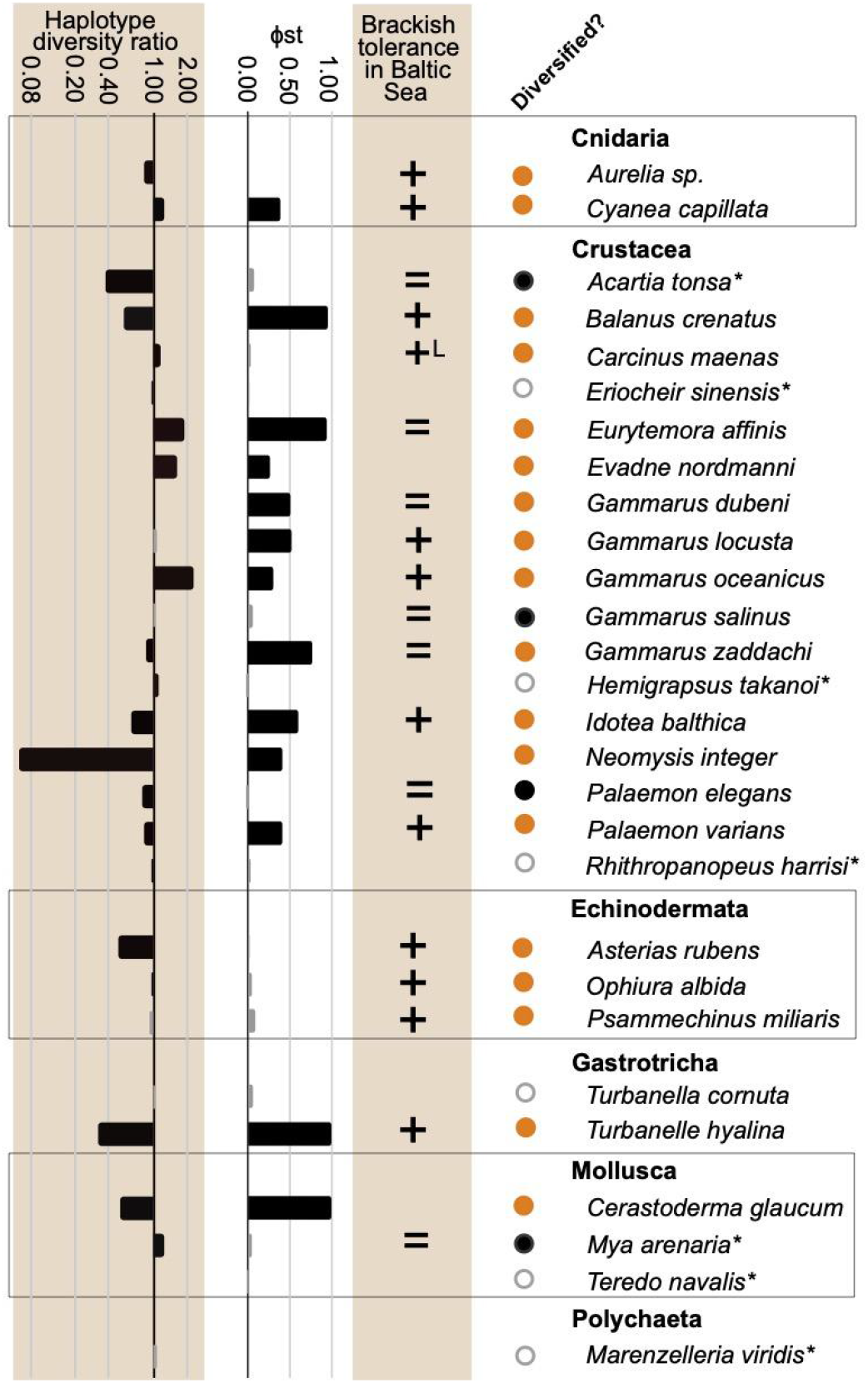
Ecological genetic summary relevant to assess the diversification potential of Baltic Sea populations. Filled orange circles indicate ecological and/or genetic diversification across the North Sea - Baltic Sea gradient, whereas filled black circles indicate the Baltic Sea population as a sink. Grey open circles represent the Baltic Sea as a putative genetic sink, but ecological confirmation is lacking. A plus indicates that the Baltic Sea population can tolerate fresher water, a superscripted “L” indicates that this was only shown for the larvae, and an equal sign indicates that there are no differences between the salinity tolerance of North and Baltic Sea populations.

### Genetic differentiation and gene flow

The three differentiation indices identified different numbers of native species as significant: Φ_ST_ was the most conservative index, differentiating 62% of native species, Jost’s D differentiated three additional natives, while Hudson’s Snn was significant for over 90%. The four native and three non-native species that were only identified by Snn had relatively small sample sizes, with the smallest population sample size per species ranging from six to nineteen, which may have caused insignificant Φ_ST_ values for some of these species. Our rarefaction analyses, however, suggested the opposite; species with small sample sizes have upward biased estimates. Alternatively, Snn overestimates genetic differentiation. Whatever the cause, these species appear not to be strongly differentiated (Fig. 4), but may be beginning to diverge. Curious are the two oldest species introductions, the soft shell clam *Mya arenaria* (ca. 1240, Petersen et al., 1992) and the Harris mud crab *Rhithropanopeus harrisii* (ca. 1870, Wolff, 2005), which had significant Snn estimates. These species may be beginning to differentiate across their genomes, which would make them ideal test cases to assess the speed of genetic and ecological diversification. Given the evolutionarily young age of the Baltic Sea, the widespread genetic differentiation between North and Baltic Sea may be surprising. However, 8000 years translate to several thousand generations for marine invertebrates, and this is more than sufficient for drift to differentiate populations, especially if those populations are small (Barton et al., 2007). Our results confirm that gene flow between the North and Baltic Sea is generally limited. The same trend is apparent for seagrass, algae, fishes, and harbour seals (Johannesson and André, 2006; Wennerström et al., 2013), suggesting ubiquitous resistance to North Sea-Baltic Sea gene-flow.

The connecting water body, the Belt Sea, is littered with islands, and, in more recent times, bridges are restricting water movement. Continuous salinity measurements show that marine water inflow from the North Sea is restricted to the cold months in most years (Lennartz et al., 2014; Ewers-Saucedo et al., 2020), which means that marine larvae, which predominantly disperse in spring and summer, will not be transported into the Baltic Sea. Our results support this limited connectivity, although it should not matter for species with highly motile adults, such as jellyfish and pelagic copepods. These species are nonetheless differentiated, which either means that very few North Sea specimens reach the Baltic Sea, or that these do not survive well under Baltic Sea conditions. Models of oceanographic connectivity for the transition zone show that locations in the Skagerrak and Kattegat and are connected and migration occurs mostly from the Kattegatt to Skagerrak (Godhe et al., 2013), but that oceanographic connectivity dropped significantly when entering the Baltic Sea (Sjöqvist et al., 2015).

Support for the adaptive hypothesis comes from three highly differentiated species (*Cerastoderma glaucum, Eurytemora affinis* and *Balanus crenatus*), for which sequence data from the transition zone between North and Baltic Sea exists. If limited dispersal is responsible for the observed pattern, the phylogeographic break between differentiated lineages should most likely be the Belt Sea. However, the Baltic Sea lineage of all three species extended past the Belt Sea, with breaks as far west as the Skagerrak. If the mitochondrial data is congruent with the rest of the genome, this points to an ecological maintenance of the two lineages, rather than a purely neutral divergence. In general, a more detailed genetic and ecological analysis of the transition zone would be highly informative.

### Non-native species and human-mediated gene flow

The differentiation index Φ_ST_ was non-significant for the seven investigated non-native species. Superficially, this result seems to counter the argument of limited connectivity between the North and Baltic Sea, as the lack of differentiation could indicate unhindered natural dispersal into the Baltic Sea. However, we consider human-mediated dispersal to play a crucial role for the colonization and differentiation processes of non-native species in the Baltic Sea (they are invasives, after all, and thus predisposed for anthropogenic dispersal). Ship traffic, particularly via the Kiel Canal, has been identified as the most likely introduction pathway for the crabs *R. harrisii* and *H. takanoi* (Nehring, 2000; Geburzi et al., 2015). It is generally considered one of the most important invasion vectors to the Baltic Sea (Leppäkoski et al., 2002; Ojaveer et al., 2017). Although not being a vector *sensu stricto*, the Kiel Canal itself provides an anthropogenic invasion corridor for species capable of long-distance migration like the Chinese mitten crab *Eriocheir sinensis*. While natural dispersal of *E. sinensis* around the Danish peninsula into the Baltic Sea would have likely taken several decades considering the dating of records from Danish coasts, it had successfully crossed the Kiel Canal west to east only six years after its first occurrence on the German North Sea coast (Herborg et al., 2003). In general, the high rate of ship traffic between the North and Baltic Seas may well cause repeated/continued introductions of non-native species that prevent differentiation of introduced Baltic Sea populations (compare Simon-Bouhet et al., 2006; Roman and Darling, 2007).

In contrast to the non-significant Φ_ST_ indices, we furthermore found the two oldest introductions, the clam *M. arenaria* and the crab *R. harrisii*, to be significantly differentiated by Jost’s D. Hudson’s Snn was even significant for all non-natives but *T. navalis* and *H. takanoi* (the latter being the most recent introduction). This may indicate the beginning of observable differentiation, and further hints at limited connectivity of the North and Baltic Sea. Alternatively, the differentiated populations were founded by different introduction events, as has been suggested for the crab *R. harrisii* (Hegele-Drywa et al., 2015). Overall, even the non-native species do not contradict the limited gene flow we observed in native species.

### Differentiation before the formation of the Baltic Sea

For four species, the cockle *Cerastoderma glaucum*, the amphipod *Gammarus locusta*, the shrimp *Palaemon varians*, and the copepod *Eurytemora affinis*, coalescent estimates dated the divergence of North and Baltic Sea populations in the Late Pleistocene between 200,000 and 450,000 years ago, often with much wider confidence intervals, but never including 8000 years. Two other species, the barnacle *Balanus crenatus* and the gastrotrich *Turbanella hyalina* show similar divergence patterns (Fig. 4), but have insufficient data to generate robust coalescent estimates. Assuming similar divergence times, Baltic Sea populations of these six species diverged from the North Sea populations much earlier than 8000 years ago. As the Baltic Sea in its current extent did not exist prior to the LGM, the divergence must have occurred somewhere else. For the cockle *C. glaucum*, long-distance dispersal from the Iberian Peninsula to the Baltic Sea aided by migrating birds has been implied based on phylogeographic reconstructions (Tarnowska et al., 2010). For the copepod *E. affinis*, this much older divergence likely dates to a previous interglacial period in today’s North Sea and East-Atlantic (Remerie et al., 2009; Winkler et al., 2011). During the Pleistocene, the British Isles were connected to the European continent with a land bridge, which separated the ancient North Sea from the southern English Channel (Cohen et al., 2017). During the subsequent glaciation of both the Baltic Sea and North Sea, the respective populations must have retreated into separate glacial refugia. The land bridge between the British Isles and Europe remained ice-free, and separated the Scandinavian and the British ice sheets due to much lower sea levels (Dawson, 1992). Marine organisms such as the amphipod *G. locusta*, the shrimp *P. varians*, the barnacle *B. crenatus* and the gastrotrich *T. hyalina* may have retreated into glacial refugia located either south of the permanent ice shields (Remerie et al., 2009; Luttikhuizen et al., 2012) or in the Irish Sea, around Scotland and in the English Channel (Roman and Palumbi, 2004; Provan et al., 2005). This means that ¼ of the native species diverged and remained separate for much longer than the current brackish water Baltic Sea has been in existence.

### Genetic diversity and population size

We found that genetic haplotype diversity and theta of Baltic Sea populations is lower in only half of the species, and nucleotide diversity in only ⅓ of the species, contrary to a previous study that found the majority of Baltic Sea populations to be less diverse (Johannesson and André, 2006). Six of those species are also significantly differentiated from the North Sea. In these species, it seems likely that the Baltic Sea populations are smaller, sustaining less genetic diversity, which is a hypothesis proposed earlier for Baltic Sea populations (Johannesson and André, 2006). The sink scenario also predicts lower genetic diversity, and genetically, this appears to be the case for six weakly differentiated species, i.e. the echinoderms *Asterias rubens* and *Ophiura albida*, the shrimp *Palaemon elegans*, and the non-natives *Rhithropanopeus harrisii, Eriocheir sinensis* and *Acartia tonsa*. However, the echinoderms are adapted to the lower salinity conditions of the Baltic Sea, which contradicts the sink scenario. For the non-natives, the sink pattern is a consequence of their recent colonization of the Baltic Sea, but does not reflect long-term conditions. In other words, not enough time has passed to identify a potential lack of gene flow after the colonization event. Moreover, they remain largely untested for rapid adaptations to the lower salinity of the Baltic Sea. Only for the shrimp *Palaemon elegans*, the Baltic Sea may represent a true sink with lower genetic diversity. That said, both Hudson’s Snn and Jost’s D differentiated the North and Baltic Sea populations of this species, which makes us wonder if this species may not have adapted to the Baltic Sea after all. The introduction of the highly divergent Black Sea lineage of *P. elegans* further complicates the issue. While we could clearly identify and exclude sequences belonging to the Black Sea lineage, it is unclear if the Atlantic lineage we considered here and the Black Sea lineage hybridize, and what the consequences may be for the Atlantic lineage.

### Life history and salinity adaptations

The fact that species with a long PLD are limited to more saline waters is intriguing. It is compatible with the fact that the larval phase is often most sensitive to environmental conditions (Sherman et al., 2016). If the planktonic larval phase is shorter, or completely absent, this may increase the probability of a species to colonize brackish to freshwater environments. This theory is well-supported by marine taxa that colonized rivers and freshwater by abbreviating or eliminating the planktonic larval phase (Vogt, 2013).

Dispersal ability did not correlate with population differentiation. This further strengthens our argument that limited water flow is not responsible for population differentiation. This mirrors results of comparative phylogeographic studies along e.g. the North and South American coasts, where dispersal ability is a poor predictor of population differentiation (Kelly and Palumbi, 2010). Instead, our results corroborate the idea that local adaptation drove population differentiation, in combination with small founding populations (Johannesson and André, 2006). Many species colonized the Baltic Sea early on, when the salinity was higher, and the connectivity to the North Sea stronger (Johannesson et al., 2011). Subsequent adaptations to declining salinities would have isolated the populations, which was exacerbated by decreasing North Sea water inflow (which is linked to the lowered salinity).

The basin-specific differences in salinity tolerance are likely due to local adaptation, driven by changes of the genomic sequence, epigenetic changes, or acclimatization. The experiments from which we derived the salinity tolerances do not allow us to distinguish between these causes, as none of them included individuals from both basins or addressed this question. Disentangling these effects will take multi-generational experiments in combination with detailed molecular approaches, but could generate unprecedented insight into the rapid evolution of freshwater tolerance. For example, common garden experiments followed by proteomics revealed several functional candidate loci that were differentially expressed between freshwater and brackish water spawning whitefish (Papakostas et al., 2012).

### What do different ecological-genetic patterns indicate?

We identified species where genetics and ecology match up, either because both suggest diversification across the North Sea - Baltic Sea gradient (7 spp.: *C. capillata, B. crenatus, G. locusta, G. oceanicus, I. balthica, P. varians, T. hyalina*), or because both suggest homogeneity (4 spp.: *A. tonsa, G. salinus, P. elegans, M. arenaria*) (Fig. 6). We also identified species with intermediate patterns, where either ecology (5 spp.: *Aurelia sp*., *C. maenas*, all 3 investigated Echinodermata spp.) or genetics alone suggest diversification (3 spp.: *E. affinis, G. duebeni, G. zaddachi*). For the remaining 9 species, comparative salinity tolerance estimates do not exist (Fig. 6).

Ecological diversification and local adaptation have been shown to occur within decades to centuries, for example cold adaptation of the invasive Burmese Python in Florida (Card et al., 2018), habitat and diet shift of Mangrove tree crabs in Georgia (Riley et al., 2014), or adaptation of mice to urban habitats (Harris et al., 2013). Thus we may expect salinity adaptation to occur rapidly and frequently, a view borne out by our data. Genetic divergence of a putatively neutral marker, such as mitochondrial DNA, occurs much slower, and only when gene flow is severely limited (Messer et al., 2016).

The diversification of ecology and genetics in concert can be attributed, on the one hand, to a much older divergence, as in *B. crenatus, T. hyalina* and *P. varians*. This means that the genome had more time to catch up with any realized lack of gene flow. The similarly strong genetic diversification in the more recently diverged amphipods *G. locusta* and *G. oceanicus*, and in the isopod *I. balthica* seems surprising considering their comparatively large population sizes (see Fig. 4, Fig. S4), as this should prevent rapid differentiation by genetic drift. However, their short generation time (Kolding and Fenchel, 1979; Leidenberger et al., 2012) might have sped up differentiation processes despite large population sizes. The lion’s mane jellyfish *C. capillata* with its highly motile adults remains a bit of a mystery in this category, which could be resolved by a phylogeographical assessment of this species.

In the investigated echinoderms, the jellyfish *Aurelia sp*. and the crab *C. maenas*, the population genetics indicate the Baltic Sea as a population sink. The ecology, however, indicates that the Baltic Sea populations are adapted to lower salinity. This could either mean that gene flow is ongoing, and they adapted in the face of gene flow (Tigano and Friesen, 2016). Alternatively, their relatively long generation time - they all need one to two years to mature (Crothers, 1967; Nichols and Barker, 1984; Jackson, 2008) - slows divergence between populations, leaving the mitochondrial genomes of these species not enough time to catch up with an actual lack of gene flow. Investigating more invertebrates with long generation times should allow us to confirm these results, for example the large conchs *Neptunea antiqua* and *Buccinum undatum*, and the hermit crab *Pagurus bernhardus*, which occur in both North and Baltic Sea (Zettler et al., 2018).

Only three species are genetically diverged but do not show signs of ecological diversification: the copepod *E. nordmanni* and the amphipods *G. duebeni* and *G. zaddachi*. These species are found deep into the Baltic Sea, and display wide salinity tolerances in both North and Baltic Sea. They indicate limited gene flow in lieu of adaptation, and further highlight that there is little connectivity into the Baltic Sea.

It is apparent that the diversity of ecological-genetic patterns in the genus *Gammarus* is highest compared to other taxonomic groups considered in our study. A potential explanation for this diversity are the different life histories of the five *Gammarus* species: While *G. duebeni, G. zaddachi* and *G. salinus* are ‘true’ brackish water species that occur in river deltas and other brackish habitats outside the Baltic Sea, *G. oceanicus* and *G. locusta* occur under (almost) fully marine conditions in the North Sea/Atlantic Ocean (den Hartog, 1964; Fenchel and Kolding, 1979; Gaston and Spicer, 2001). The latter two species therefore had to evolve wider salinity tolerances to inhabit the Baltic Sea. As a consequence of their comparatively long divergence time (this paper for *G*. *locusta*, Normant et al., 2005 for *G*. *oceanicus*), their ecological diversification is reflected by neutral genetic differentiation. For *G. duebeni* and *G. zaddachi*, on the other hand, the genetic divergence might be reflected by diversification on an ecological trait other than salinity tolerance. In fact, Kolding and Fenchel (1979) found differing reproductive traits in North and Baltic Sea populations of both species. In contrast to the diversity in *Gammarus*, the congruent pattern we found in all three echinoderms might be a consequence of their common evolutionary origin as fully marine species, leading to similar adaptation processes to lower salinities during their colonization of the Baltic Sea.

For the three species fitting the sink scenario (*A. tonsa, G. salinus* and *P. elegans*), their concordant pattern of homogeneity at the genetic and ecological level could be explained by individuals being swept into the Baltic Sea without forming reproducing populations. However, given the lack of gene flow between North and Baltic Sea we and others observed for the majority of organisms (Johannesson and André, 2006; Sjöqvist et al., 2015) in combination with the limited oceanographic connectivity that has been modeled (Barz et al., 2006; Hordoir et al., 2013), this scenario appears unlikely to us. Moreover, these species are common in the Baltic Sea, and not limited to the most western parts. Instead, this apparent homogeneity is attributable on the one hand to a recent colonization of the Baltic Sea of species with a wide salinity tolerance, i.e. the non-natives *A. tonsa* and *M. arenaria*. This may also be the case for the amphipod *G. salinus*, which we consider the only native species that clearly fits the sink scenario. On the other hand, the homogeneity might be attributable to a beginning divergence, as we assume for the shrimp *P. elegans*, which we found significantly differentiated by two of the differentiation indices, but not by Φ_ST_. Furthermore, the recent introduction of the highly divergent Black Sea lineage of *P. elegans* may complicate the assessment of salinity tolerance. This species clearly warrants further investigation, but for the moment, the Baltic Sea may be considered a sink for this species.

In summary, our data provide evidence for the coexistence of divergent eco-evolutionary trajectories in different marine invertebrate species that inhabit the North Sea - Baltic Sea region. These trajectories appear to be shaped by a complex interplay of the species’ ecology, evolutionary background, and colonization history (compare Ewers-Saucedo and Wares, 2020). Future studies including both, additional life-history traits and genomic/non-neutral genetic markers could draw a more detailed picture of the formation of these trajectories.

### The Baltic Sea: a natural experiment on the tempo and mode of adaptation

We show that the North Sea - Baltic Sea environmental gradient represents a diversification hotspot for most marine invertebrates, many of which have altered salinity tolerances in the Baltic Sea. This genetic and ecological divergence is congruent with similar studies on fishes and algae (Johannesson and André, 2006; Papakostas et al., 2012; Wennerström et al., 2013; Berg et al., 2015; Guo et al., 2015; Sjöqvist et al., 2015), which makes this gradient ubiquitous across phyla. The Baltic Sea provides the unique opportunity to understand the mechanisms underlying this diversification, and particularly the mechanisms of salinity adaptation. Especially the combined use of non-native and native species would allow a comparison of species at different stages of a colonization process that is almost certain to require adaptation; from ongoing colonization and expansion as in the crab *H. takanoi*, to recent colonization within the last few centuries as in in the mitten crab *E. sinensis*, or colonization many centuries ago as in the clam *M. arenaria* to several thousand years as in the amphipod *G. duebeni* and finally a divergence several hundred thousand years as in the amphipod *G. locusta*. The physical proximity between the North and Baltic Sea facilitates ecological common garden experiments across a natural salinity gradient (Sjöqvist et al., 2015). With the emerging suite of genomic tools, such experiments advance our understanding of adaptation and colonization (Sherman et al., 2016). While they have been predominantly restricted to fishes (Larsen et al., 2008; Papakostas et al., 2012), our results highlight the great potential of transferring such approaches to further (invertebrate) species to achieve a more complete understanding of the evolution of the Baltc Sea’s unique species community.

## Acknowledgements

We would like to thank Andrea Böttcher, Selena Djabri, Andreas Dürr, Ursula Maasz, Hannah Meenke, Nena Weiler, and Marina Wirth for their data contributions, and Prof. Dr. Günter Hartl for supporting data generation. We appreciate the data sent to us by Prof. Dr. Christoph Schubart and Dr. Silke Reuschel. This study was financially supported by a grant from the German Federal Ministry of Education and Research (Bundesministerium für Bildung und Forschung, project number 01UQ1711).

