## Supporting Information 3 for "A marginal habitat, but not a sink: Ecological genetics reveal a diversification hotspot for marine invertebrates in the brackish Baltic Sea"

**Supporting Information 3 – Supplemental figures**

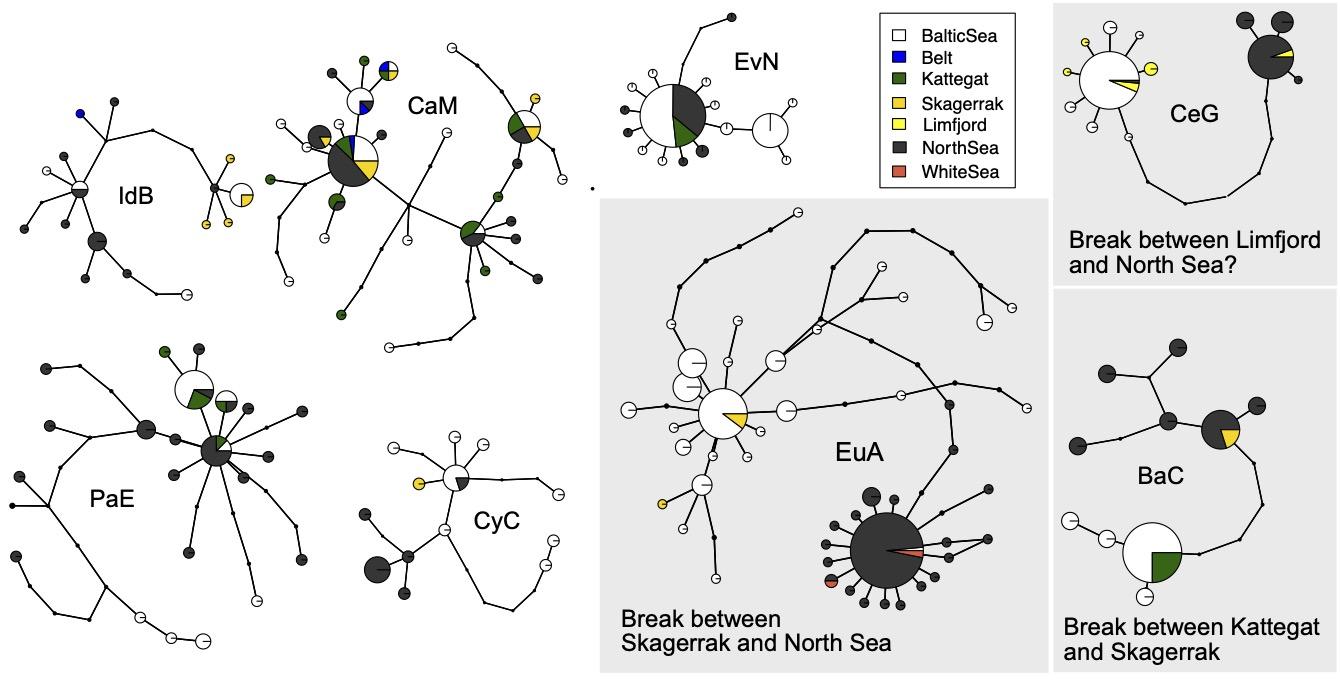

**Figure S3.1.** Extended haplotype networks for all species, for which sequences from the transition zone (as defined in Fig. 1 of the main manuscript) existed. The grey boxes indicate species with high differentiation between North and Baltic Sea populations, which allowed the inference of the phylogeographic break between these lineages. Abbreviations as in Tab. 1 of the main article.

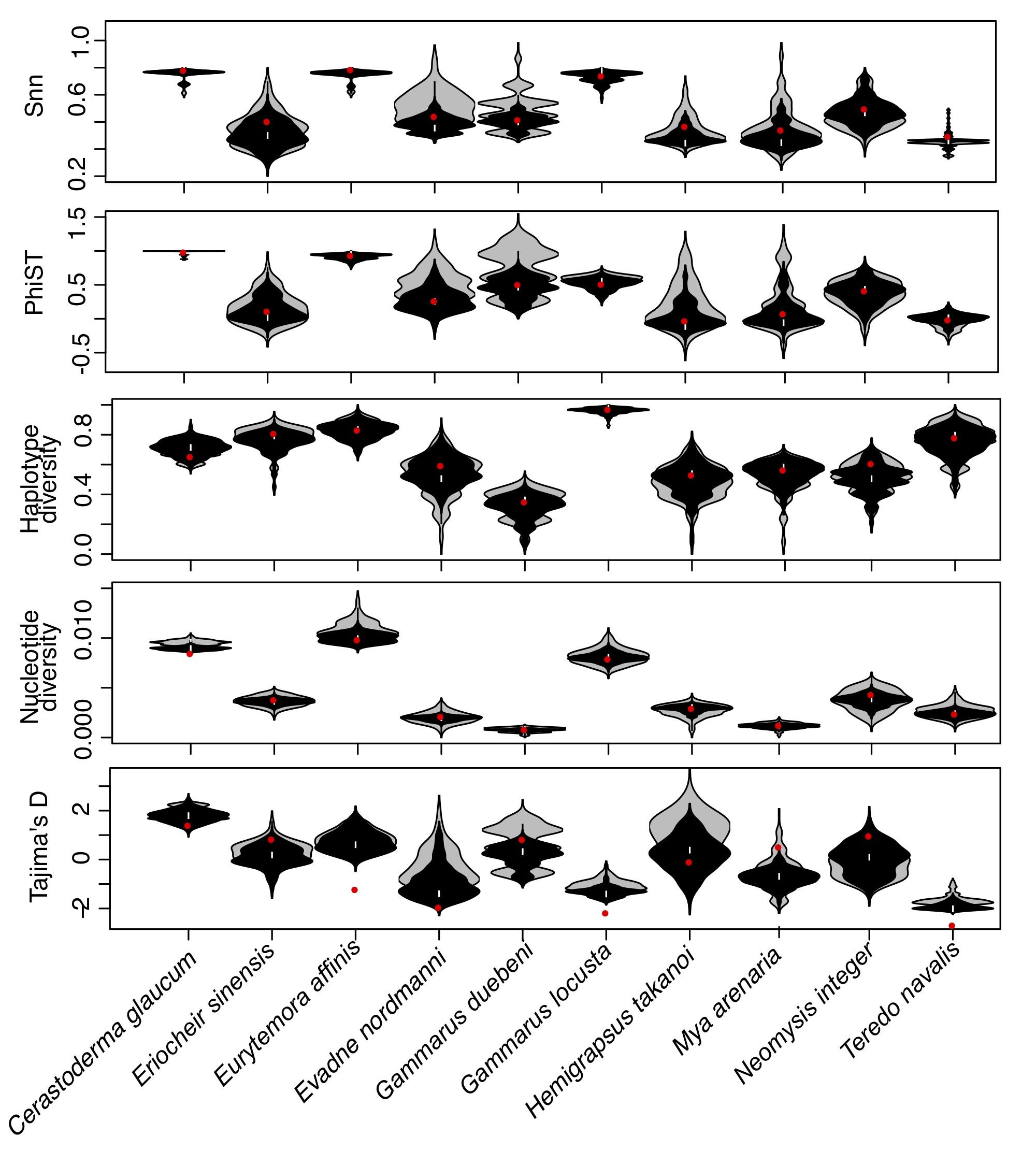

**Figure S3.2.** Rarefaction analysis. Distribution of population statistic estimates across 100 random subsamples taken from species with large population sample sizes. Grey violin plots are based on five sequences per population, and black violin plots on 7 sequences. Red dot indicates estimate based on all available (> 20) sequences.

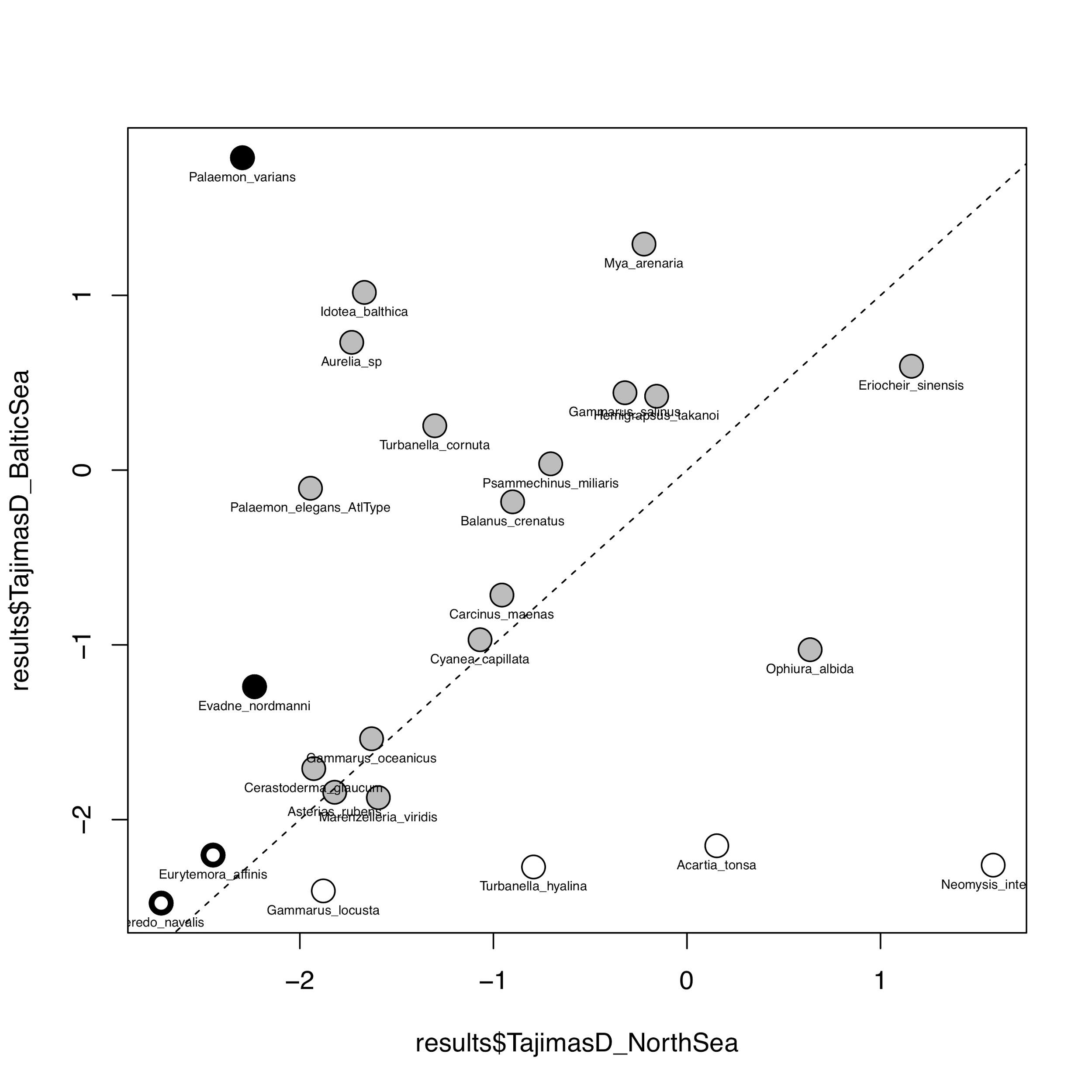

**Figure S3.3.** Estimates of Tajima’s D. Grey: Tajima’s D not significantly different from zero for either population, black: Tajima’s D significantly different from zero in the North Sea population, white: Tajima’s D significantly different from zero in the Baltic Sea population, and thick black circles: both populations significantly different from zero.

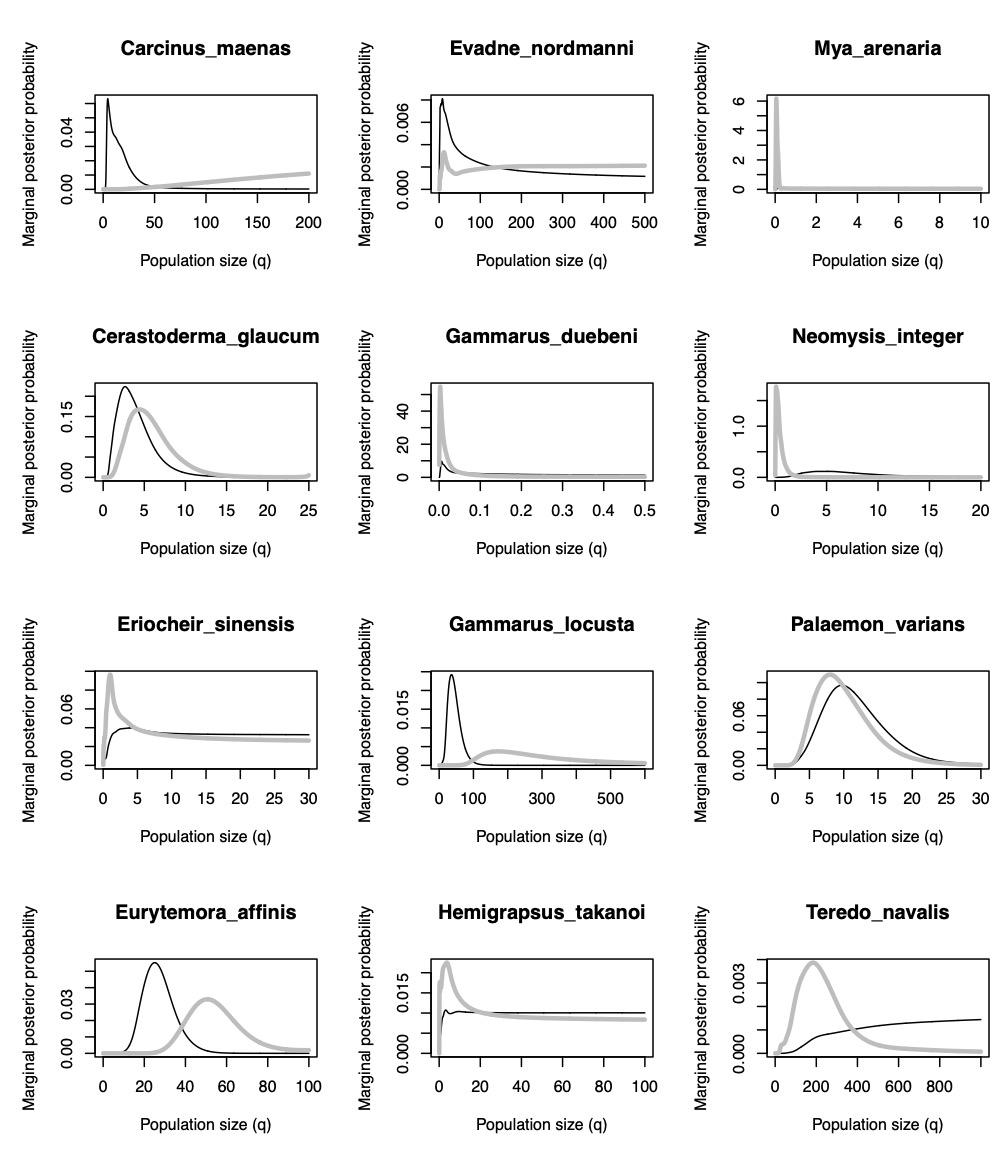
**Figure S3.4.** Coalescent estimates of mutation-rate scaled population size q obtained with IMa2. Grey= Baltic Sea, black = North Sea.

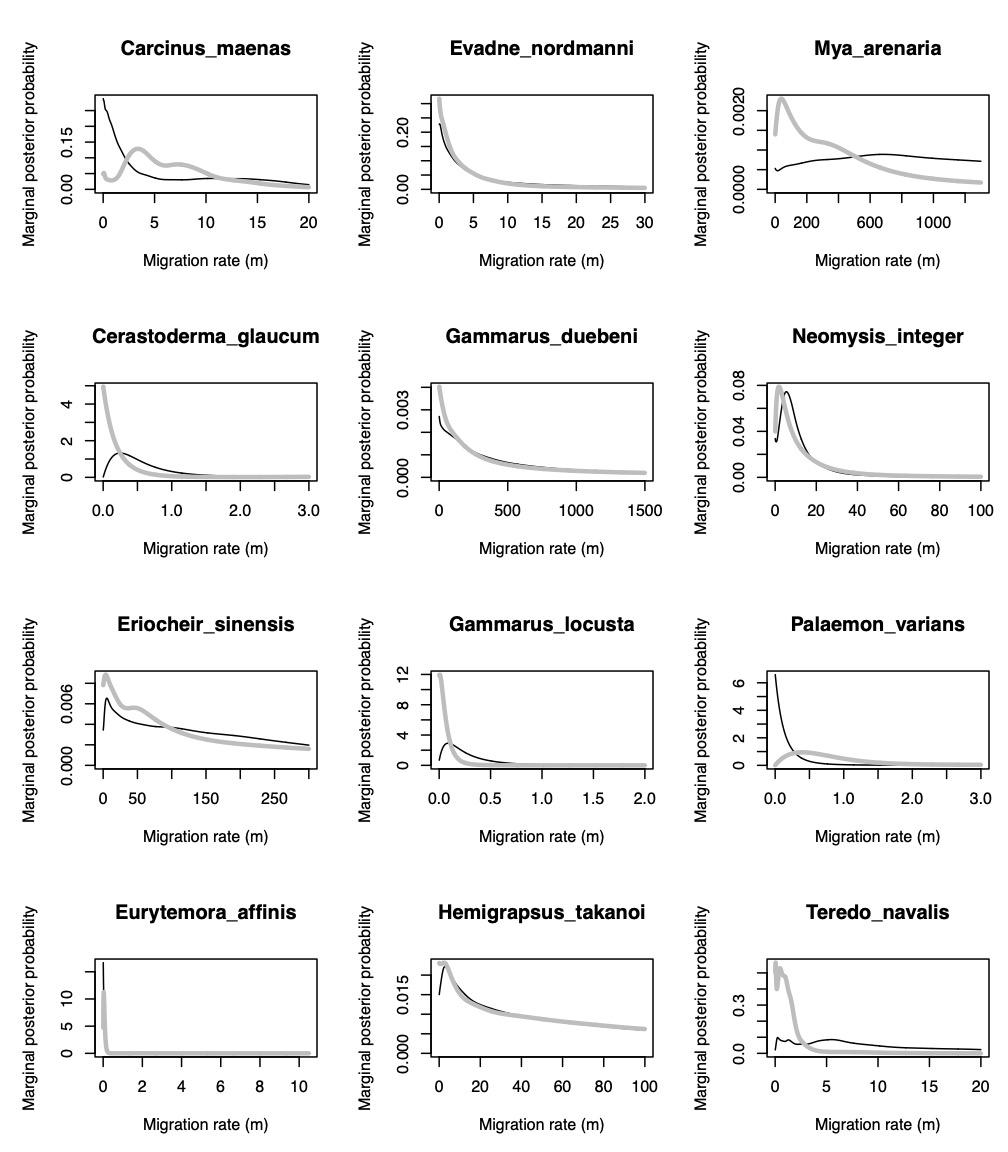

**Figure S3.5.** Coalescent estimates of mutation rate-scaled migration rates m obtained with IMa2. Grey = migration rate from Baltic Sea to North Sea, black = migration rate from North Sea to Baltic Sea.
